# Targeting Langerhans cells using a modular mannosylated nucleic acid-based vaccine platform

**DOI:** 10.1101/2025.06.04.657560

**Authors:** Simon Christian Vinther, Antonia Resag, Karina Thao Thu Le, Claudia Christine Zelle-Rieser, Helen Strandt, Mette Galsgaard Malle, Stig Hill Christiansen, Barbara Del Frari, Theresia Stigger, Bastian Schilling, Jonas Wohlfarth, Steffen Thiel, Julián Valero, Patrizia Stoitzner, Jørgen Kjems

## Abstract

The skin is a diverse reservoir of immune cells with strong potential for immunotherapeutic delivery. Langerhans cells (LCs) in the epidermis are antigen-presenting cells, accessible for vaccination using carbohydrate-conjugated therapeutics targeting their endocytic lectin receptor, Langerin. As carbohydrate-lectin binding is highly dependent on valency, scaffolds that enable control over ligand spacing and stoichiometry are instrumental in enhancing receptor binding and selectivity. Here we utilized a self-assembled nucleic acid-based Holliday Junction scaffold, fully modified for nuclease protection and with a well-defined carbohydrate arrangement to optimize drug delivery to LCs. In vitro screening with Langerin-expressing cells revealed that mannosylated Holliday Junctions showed the strongest binding. This was confirmed in human epidermal cell suspensions, demonstrating specificity and valency-driven interactions. Topical administration of mannosylated scaffolds on skin explants enabled effective targeting of epidermal LCs. Finally, in an antigen-presentation assay, in vitro differentiated LCs loaded with mannosylated and peptide-conjugated scaffolds significantly enhanced T cell activation. Overall, our study presents a promising nucleic acid-based platform for precise LC targeting and drug delivery, with broad potential for skin-directed immunotherapies.

## Introduction

Targeted drug delivery has been revolutionized by leveraging endocytic C-type lectin receptors (CLRs) using carbohydrate-conjugated therapeutics^1,2^. The carbohydrates can interact with CLRs through binding to a conserved carbohydrate-recognition domain (CRD) in a Ca^2+^-dependent or independent manner^3^. In mammalian CLRs, monosaccharides only transiently bind to the CRD due to their low mM-range affinity. However, stronger binding interactions can be achieved using multivalent carbohydrate ligands to induce avidity, known as the glycocluster effect^4,5^, whilst potentially enhancing receptor selectivity^6^.

Carbohydrate-conjugated therapeutics have also been employed for targeting dendritic cells (DCs), which, as sentinels of the immune systems, serve as entry points for antigen loading in situ^7,8^. This approach effectively leads to antigen presentation to T cells by the DCs and promotes robust immunity to infection and cancer ^8,9^. In particular, the dermal region in the skin is a rich source of tissue-resident DCs and macrophages, expressing carbohydrate-binding CLRs such as DC-specific intercellular adhesion molecule-3-grabbing non-integrin (DC-SIGN) and the mannose receptor (MR)^7,8,10^. Another unique type of antigen-presenting cell within the outer epidermal skin layers is Langerhans cells (LCs)^11,12^. They express Langerin, a carbohydrate-binding CLR with a Ca^2+^-dependent CRD which upon binding of ligands leads to internalization into specialized intracellular compartments (Birbeck granules)^13,14^. Prior investigations using antibody-conjugated antigens have demonstrated the ability of LCs to present internalized antigens via Langerin to both CD4^+^ and CD8^+^ T cells^15–17^. Langerin targeting can also be achieved using appropriate carbohydrates. After endosomal uptake, the carbohydrates are released upon acidification, ensuring the receptor does not facilitate expulsion of the conjugated antigen through recycling mechanisms^18^. Langerin contains a glutamate-proline-asparagine (EPN) domain which is highly conserved in the CRD of numerous CLRs, facilitating affinity and selectivity for carbohydrates containing terminal mannose (Man), fucose (Fuc), glucose (Glu), and N-acetyl glucosamine (GlcNAc)^3,19^. In recent years, chemically modified carbohydrates, known as glycomimetics, have also been developed to enhance Langerin-specific LC targeting^20,21^. Carbohydrate-conjugates for Langerin targeting have previously been used on peptide-^22–24^, polymer-^25,26^ and liposome-based scaffolds^24^, offering a varying degree of ligand control, targeting efficacy and uptake potential for antigen delivery. Despite significant advancements, these technologies have some drawbacks resulting in heterogeneous structures and poor control over ligand arrangement, potentially affecting the avidity-based Langerin targeting, as well as instability and immunogenicity when used in vivo^27,28^. Instead, using biomaterial as a well-defined uniform scaffold for controlled ligand positioning may be favored to enhance Langerin targeting.

This is achievable with nucleic acids-based scaffolds, which can be designed to self-assemble in a sequence-defined manner following Watson-Crick base pairing^29,30^. The incorporation of ribose-modified nucleotides into nucleic acid-based scaffolds such as 2’-fluoro (2’-F)^31^, 2’-O-methyl (2’-OMe)^32^ and 2’-O,4’-C-methylene-bridges in locked nucleic acids (LNA)^33,34^ enhances their stability against nuclease degradation and reduces unwanted immune effects, enabling their application in vivo ^35^. Nucleic acid-based scaffolds can also be functionalized on individual strands prior to assembly, enabling the construction of drug delivery vehicles with nanometer control of the targeting ligand position, geometry and stoichiometry, whilst a variety of therapeutic payload can be conjugated to different strands^36,37^. Despite these unique characteristics, a glycosylated nucleic acid-based scaffold has to our knowledge not yet been employed for targeted drug delivery to LCs.

In this study we designed a carbohydrate-conjugated nucleic acid-based scaffold, for which ligand valency was optimized for efficient binding to Langerin and subsequent intracellular uptake. We demonstrate that the scaffold could specifically target human LCs and efficiently deliver antigens for T cell presentation, demonstrating the potential of our novel nucleic acid-based vaccine platform for future skin immunotherapy.

## Results

### Carbohydrate screening demonstrates Langerin preference for Man

A previously characterized 2’-OMe and LNA-modified Holliday Junction (HJ) was used as a scaffold for displaying targeting ligands, due to its robust self-assembly and extraordinary in vivo stability^38^. The scaffold is comprised of four 12-nucleotide long strands (Q1 – Q4), each with a 5’ C6-amino linker for chemical functionalization, which self-assembles stoichiometrically upon equimolar mixing (Figure 1A). The HJ enables modular assembly of biomolecules by conjugating each individual oligonucleotide strand before mixing, providing an extended pharmacokinetic profile and precise biomolecular targeting into a single scaffold, previously employed for developing hepatic- or tumor-targeting nanomedicines^38–40^. In the interest of targeting Langerin on epidermal LCs, the HJ scaffold was employed with a carbohydrate ligand display with different valencies.

**Figure 1.**
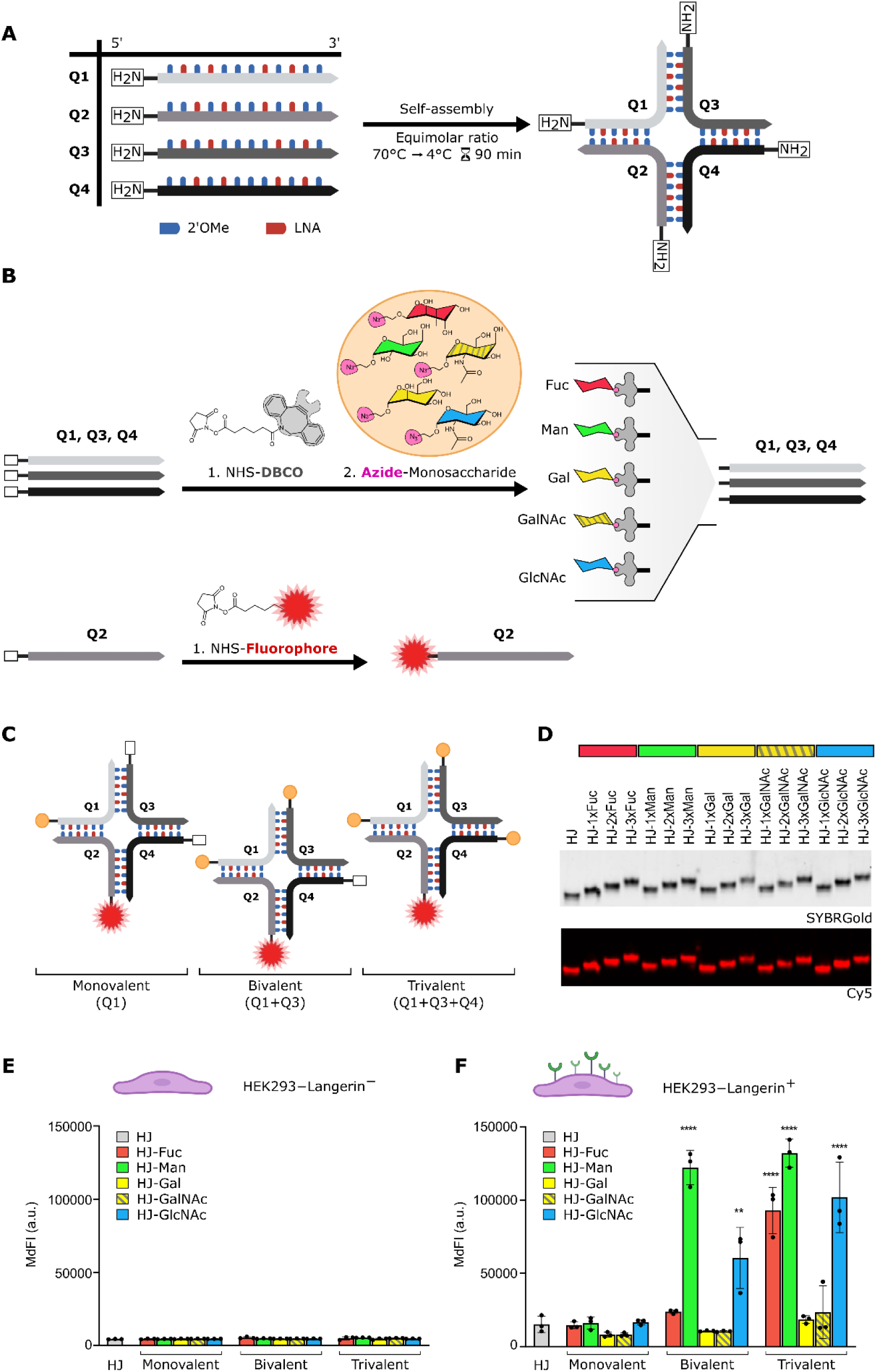
Assembly of glycosylated HJ scaffolds for screening binding to Langerin-expressing HEK293 cells. **A**) Schematic illustration showing the self-assembled HJ scaffold comprised of four strands (Q1-Q4) with 2’OMe (*blue)* and LNA (*red)* modifications, and 5’-amino functionalization (*white box*). **B**) Reaction scheme for functionalization of Q1, Q3 and Q4 with NHS-DBCO (**1**), followed by a SPAAC reaction to azide-modified monosaccharides (**2**) (color indexed: Fuc = *red,* Man = *green,* Gal = *yellow*, GalNAc = *striated-yellow*, GlcNAc = *blue*). Q2 was labeled with NHS-modified fluorophores (**1**). **C**) Schematic representation of assembled glycosylated (*orange)* HJs in a mono, bi- or trivalent display with a Cy5-labeled Q2. **D**) 16 % native PAGE visualizing assembled glycosylated HJs with SYBRGold (*top*) and Cy5 signal (*bottom*). **E**) Binding of glycosylated HJs to HEK293-Langerin^−^ and HEK293-Langerin^+^ cells as measured by the median fluorescent intensity (MdFI) of the Cy5 fluorescence signal on HJs using flow cytometry. Results shown from n = 3 technical replicates and expressed by the mean ± SD. Statistical analysis was conducted by a one-way ANOVA in comparison to the non-glycosylated HJ scaffold: p-value * = <0.05, ** = <0.01, *** = <0.001, ****= <0.0001.

To investigate the optimal carbohydrate ligand composition for targeting Langerin using the HJ scaffold, five different monosaccharides were individually conjugated to each strand. The Q1, Q3, and Q4 strands were initially DBCO-labeled by NHS-amine reactivity, followed by a strain-promoted azide-alkyne cycloaddition (SPAAC) reaction to azide-functionalized Fuc, Man, galactose (Gal), N-acetyl galactosamine (GalNAc) and GlcNAc (Figure 1B). The Q2 strand was reacted with NHS-ester functionalized fluorophores to enable detection of HJ scaffolds by flow cytometry and confocal microscopy in subsequent experiments. The functionalized Q1-Q4 strands were purified by reverse-phase high-performance liquid chromatography (RP-HPLC) and validated by denaturing polyacrylamide gel electrophoresis (PAGE) (Supplementary Figure S1).

The glycosylated HJs were arranged with carbohydrate-conjugated strands in a monovalent display on Q1, bivalent on Q1+Q3, or trivalent on Q1+Q3+Q4, each with a Cy5-labeled Q2 (Figure 1C). Following heat annealing, the formation of glycosylated HJs was successfully confirmed by native PAGE (Figure 1D).

To evaluate binding of glycosylated HJs to cells, a screening assay was conducted with Langerin-expressing HEK293 cells (HEK293-Langerin^+^). The cells were incubated with 50 nM of Cy5-labeled glycosylated HJs for 30 min prior to analysis by flow cytometry. Binding specificity of the glycosylated HJ scaffolds was validated on cells without Langerin expression (HEK293-Langerin^−^) and compared to the non-glycosylated HJ, which showed no increase in median fluorescent intensity (MdFI) (Figure 1E). In contrast, HEK293-Langerin^+^ cells showed significant binding of HJ-3xFuc and both bi- and trivalent HJ-GlcNAc (HJ-2xGlcNAc, HJ-3xGlcNAc) and HJ-Man (HJ-2xMan, HJ-3xMan), compared to the non-glycosylated HJ control (Figure 1F). No binding was observed for HJ-Gal and HJ-GalNAc scaffolds. Interestingly, a stepwise binding enhancement correlating with the number of ligands was observed for HJ-GlcNAc, while HJ-Man showed a 7.6-fold increase from HJ-1xMan to HJ-2xMan, but only a marginal 1.1-fold increase from HJ-2xMan to HJ-3xMan. The highest binding was obtained for HJ-3xMan, although not statistical difference from HJ-3xGlcNAc was measured.

Together, an overall higher binding of HJ-Man scaffolds was observed compared to other monosaccharides, suggesting a preference for Langerin binding to mannosylated HJ scaffolds.

### Enhanced binding to Langerin-expressing cells with a trimannose ligand

Increasing the carbohydrate ligand valency often results in stronger binding, which is consistent with the reported glycocluster effect for CLR targeting ^23,41^. Therefore, the Langerin binding properties of a trimannose (TriMan) ligand on the HJ scaffold were explored. Q1, Q3 and Q4 strands were labeled with an alkyne-functionalized branched linker by NHS-amine reactivity, followed by copper-catalyzed azide-alkyne cycloaddition (CuAAC) reaction to azide-functionalized Man (Figure 2A). TriMan-labeled strands were purified by RP-HPLC and validated by denaturing PAGE (Supplementary Figure S2).

**Figure 2.**
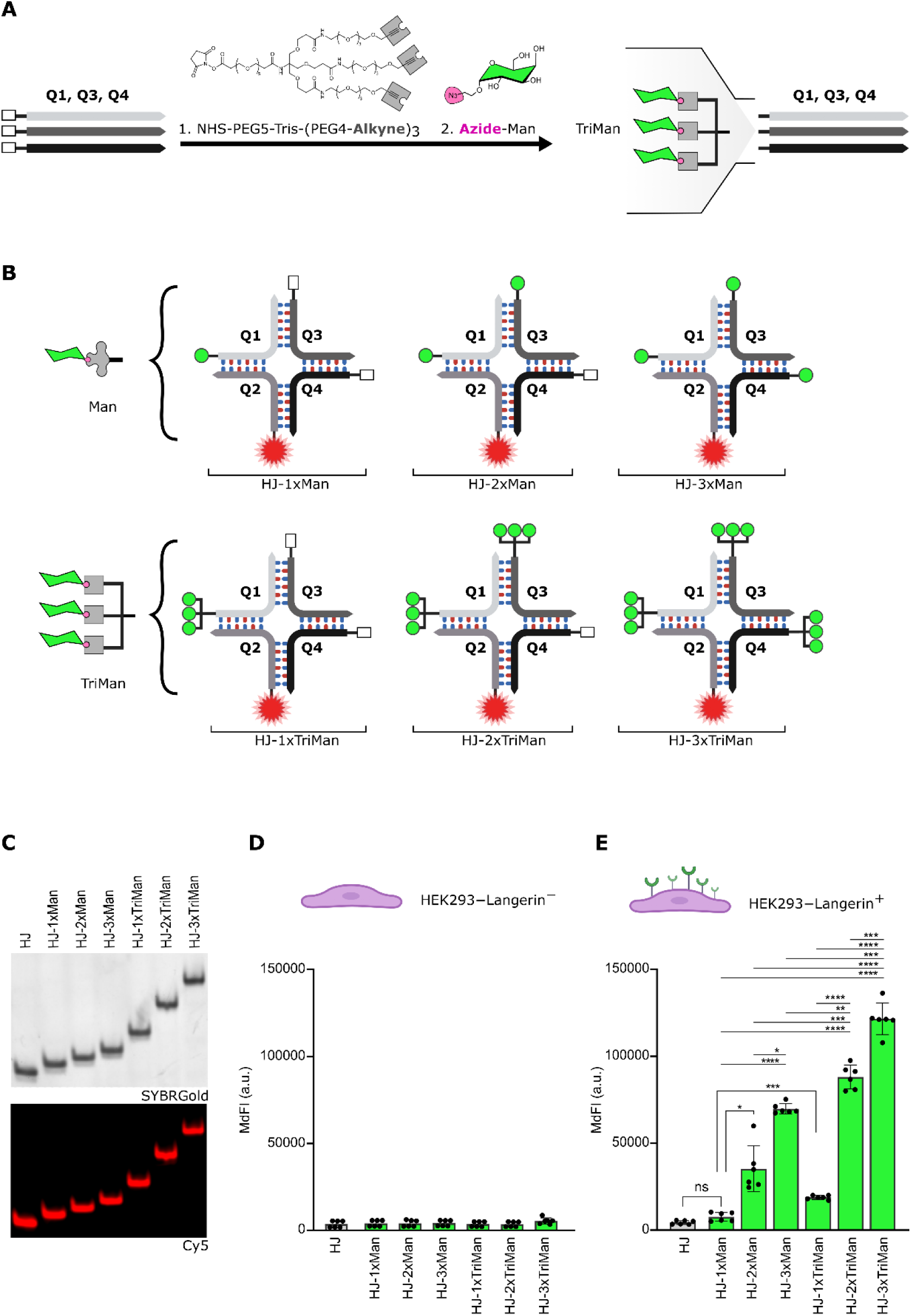
TriMan synthesis, HJ conjugation and binding characterization to Langerin-expressing HEK293 cells. **A**) Reaction scheme for TriMan-functionalization of Q1, Q3 and Q4 with an alkyne-modified branched linker, NHS-PEG5-Tris-(PEG4-Alkyne)_3_ (**1**), followed by a CuAAC reaction to azide-modified Man (**2**). **B**) Schematic representation of Man or TriMan-functionalized HJs in a mono-, bi or trivalent display with a Cy5-labeled Q2. **C**) 16 % native PAGE visualizing the assembled mannosylated HJs from SYBRGold (*top*) and Cy5 signal (*bottom*). **D-E**) Binding of mannosylated HJs to HEK293-Langerin^−^ and HEK293-Langerin^+^ cells as measured by MdFI of the Cy5 fluorescence signal on HJs using flow cytometry. Results shown from n = 3 technical replicates from two independent experiments and expressed by the mean ± SD. Statistical analysis was conducted by a one-way ANOVA followed by a Tukey’s post hoc test. p-value * = <0.05, ** = <0.01, *** = <0.001, ****= <0.0001.

Mannosylated HJs were assembled with Man- or TriMan-modified strands as previously described in a mono- (Q1), bi- (Q1+Q3), and trivalent (Q1+Q3+Q4) arrangement with a Cy5-labeled Q2 (Figure 2B), forming uniform scaffolds as validated by native PAGE (Figure 2C).

Considering that binding to HEK293-Langerin^+^ cells during the previous monosaccharide screening assay was stagnant between HJ-2xMan and HJ-3xMan at 50 nM, the concentration was reduced to 25 nM. The lower concentration circumvented saturation and yielded valency-dependent binding of Man- and TriMan-conjugated HJs to HEK293-Langerin^+^ cells (Figure 2D-E). The ligand enhancement from HJ-1xMan to HJ-3xMan increased the relative binding 9-fold, and for HJ-1xTriMan to HJ-3xTriMan 6.6-fold. Furthermore, within all HJ ligand arrangements (mono, bi and trivalent display), the TriMan ligand significantly enhanced binding to HEK293-Langerin^+^ cells by 1.8-2.5-fold compared to similar arrangements with the Man ligand. Interestingly, comparing HJ-3xMan and HJ-1xTriMan with equimolar mannosylation revealed a 3-fold binding reduction of the latter, suggesting a favored spatial orientation of ligands on the HJ scaffold in a bivalent (Q1+Q3) or trivalent (Q1+Q3+Q4) display for Langerin binding.

Different arrangements for mono- and bivalent mannosylated HJs revealed a 1.2-fold binding enhancement for the bivalent Q1+Q3 configuration compared to Q1+Q4 and Q3+Q4 (Supplementary Figure S3). These results highlight the importance of ligand orientation for Langerin targeting using the HJ scaffold, therefore the bivalent Q1+Q3 configuration was used for HJ-2xMan and HJ-2xTriMan in subsequent experiments.

In summary, employing a TriMan ligand in a bi- or trivalent arrangement on the HJ scaffold may improve Langerin binding and facilitate more effective targeting of LCs.

### Mannosylated HJs are engulfed via Langerin and accumulate in lysosomal compartments

The endocytic Langerin receptor on LCs can be harnessed for trafficking carbohydrate-conjugated antigens into endo-lysosomal compartments for processing and presentation to T cells^22^. For this reason, Langerin-mediated uptake of mannosylated HJs was investigated in HEK293-Langerin^+^ cells by removing non-internalized scaffolds. This was achieved using an EDTA-supplemented wash buffer for chelating Ca^2+^, thereby disrupting the Ca^2+^-dependent binding of mannosylated HJs to the extracellular CRD (Supplementary Figure S4). Implementing these washing steps after incubation of Cy5-labeled mannosylated HJs with HEK293-Langerin^+^ cells for up to 300 min allowed uptake to be characterized by flow cytometry (Figure 3A). After 60 min of incubation, an increase in MdFI indicated an emerging uptake of all mannosylated HJ scaffold, whereas the non-glycosylated HJ remained stagnant. After 300 min, HJ-3xMan, and more prominently HJ-3xTriMan, exhibited the highest cellular uptake.

**Figure 3.**
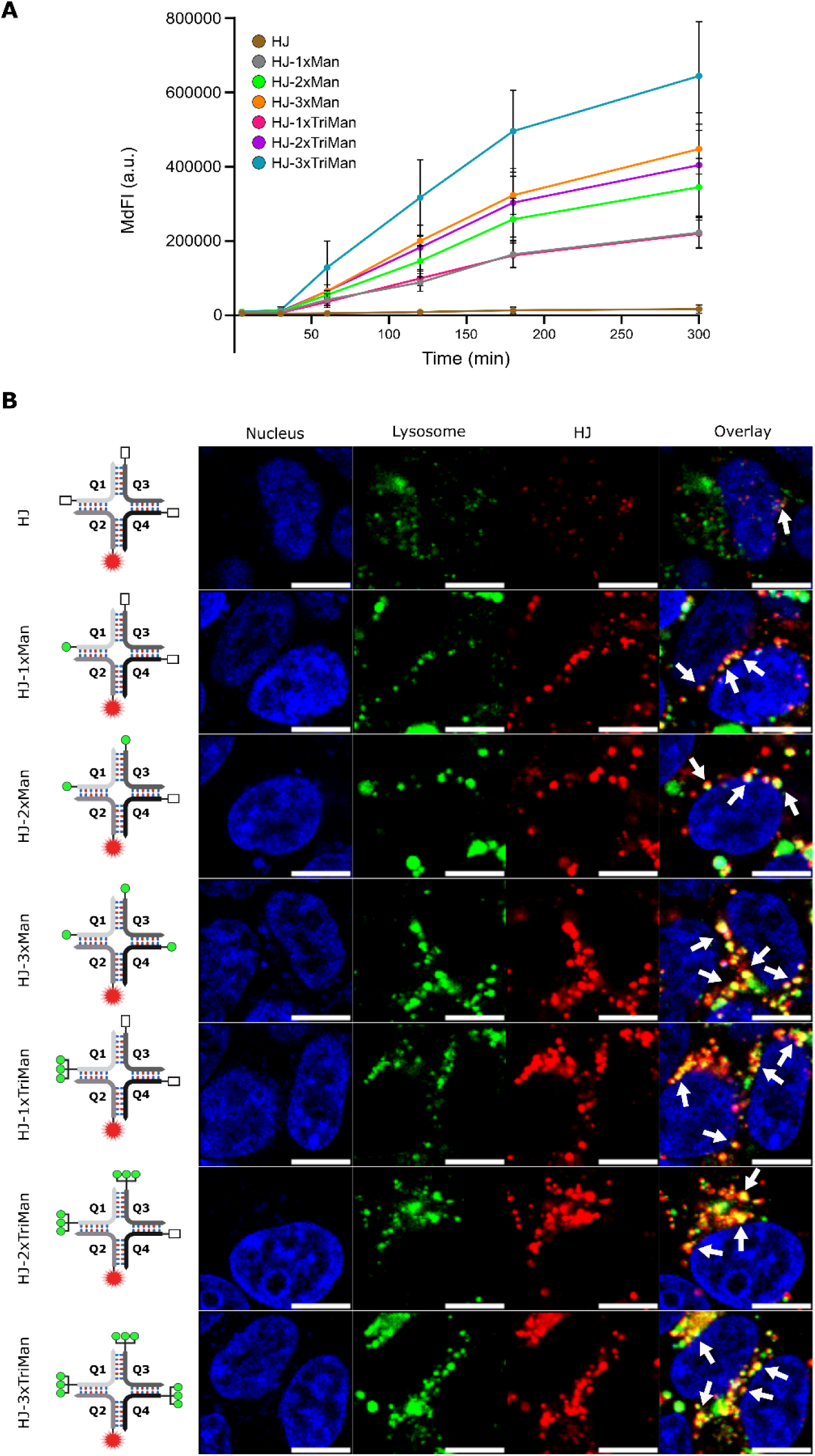
Uptake and lysosomal accumulation of differently mannosylated HJs into Langerin-expressing HEK293 cells. **A**) Kinetics for uptake of Cy5-labeled mannosylated HJs in HEK293-Langerin^+^ cells over 5 – 300 min at 37°C. MdFI results are shown for three independent experiments and expressed by the mean ± SD. **B**) Representative confocal microscopy images of HEK293-Langerin^+^ cells incubated with Cy5-labeled, mannosylated HJ scaffolds (*red*). Cells were stained with LysoTracker^TM^ DND-26 green (*green*) and nucleus stained with Hoechst 33342 (*blue*). Arrows indicate colocalization between mannosylated HJ scaffolds and lysosomal compartments. Scale bar: 10 µm.

Intracellular trafficking of mannosylated HJs was characterized by incubation of HEK293-Langerin^+^ cells overnight with a lysosomal dye (LysoTracker^TM^ DND-26 green). Visualization by live-cell microscopy confirmed the expected negligible uptake of non-glycosylated HJ and the efficient internalization of mannosylated HJs into confined cellular compartments, which colocalized with the lysosomal dye (Figure 3B).

These results demonstrate the effective Langerin-mediated uptake and trafficking of mannosylated HJs to lysosomal compartments, enabling their use for antigen delivery and processing.

### Specific targeting of mannosylated HJs to LCs in human epidermal cell suspensions

Following the promising characterization of mannosylated HJs binding and uptake in a Langerin-transfected cell line in vitro, targeting specificity of human LCs within epidermal cell suspensions was determined ex vivo.

For this purpose, epidermal cell suspensions were prepared from human skin samples and incubated overnight with 20 nM ATTO647N-labeled mannosylated HJs. To evaluate the targeting specificity of mannosylated HJs to epidermal cells, flow cytometry was performed using antibodies to detect LCs (CD45^+^, HLA-DR^+^, CD1a^+^) and to separate them from keratinocytes (CD45^−^) (Figure 4A-C). Donor-to-donor variations were addressed by normalizing to the non-glycosylated HJ-treated sample from the same donor.

**Figure 4.**
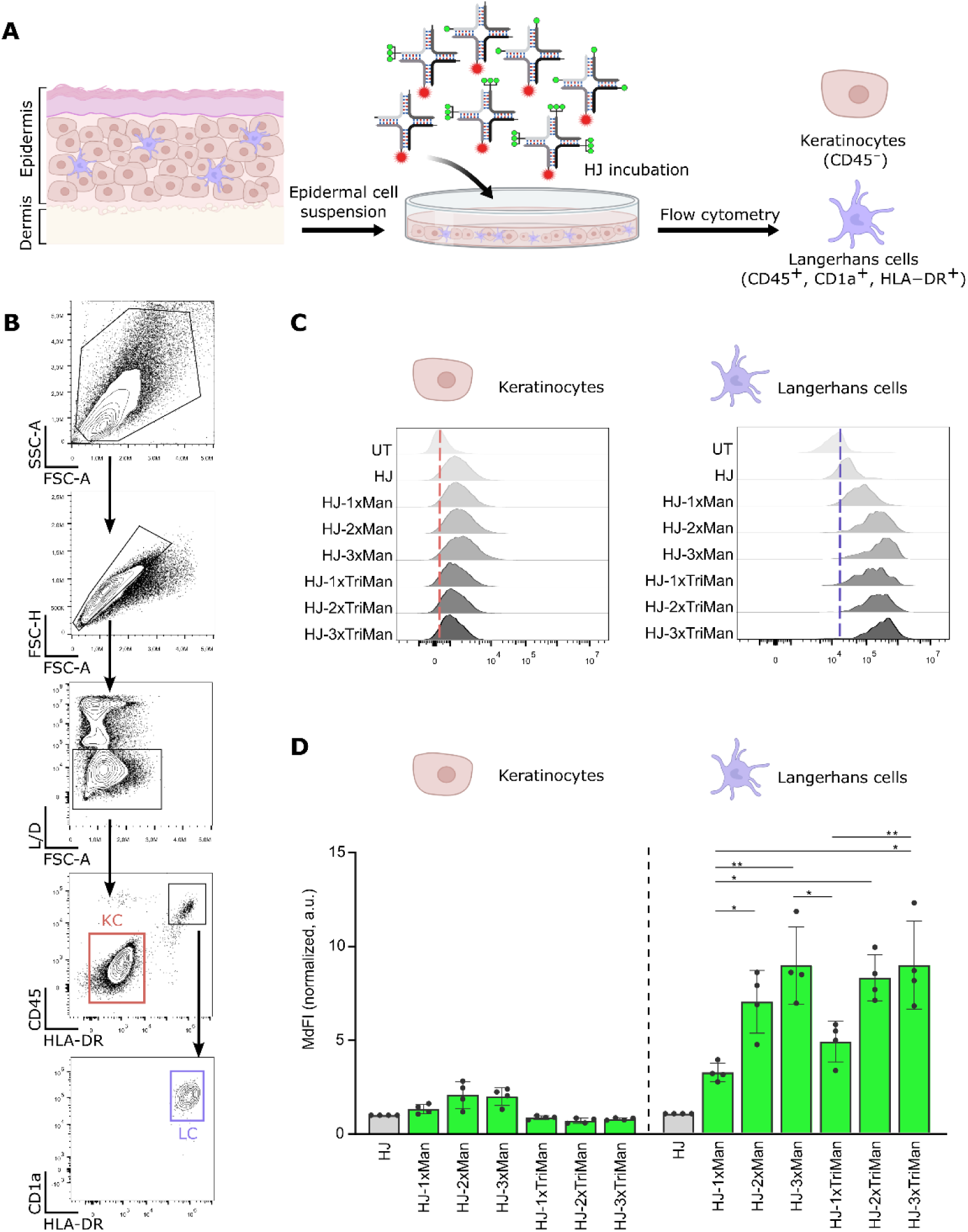
Targeting LCs in human epidermal cell suspensions with mannosylated HJs. **A**) Visual representation of the preparation of epidermal cell suspensions from human donors, followed by incubation with fluorescently labeled, mannosylated HJs and flow cytometry analysis of LCs and keratinocytes. **B**) Gating strategy demonstrating keratinocytes (KC, *red*) and LCs (LC, *violet*) in epidermal cell suspensions, identified as viable CD45^−^ or CD45^+^, HLA-DR^+^, CD1a^+^ cell populations, respectively. **C**) Representative histograms are shown for the fluorescent signal of ATTO647N-labeled HJ scaffolds in keratinocytes and LCs. **D**) Normalized MdFI of the ATTO647N fluorescent signal conjugated to the HJ scaffold in the keratinocyte and LC populations from four independent human donors. Results expressed by the mean ± SD of MdFI normalized within each donor to the non-glycosylated HJ. Statistical analysis was conducted by a one-way ANOVA followed by a Tukey’s post hoc test. p-value * = <0.05, ** = <0.01, *** = <0.001, ****= <0.0001.

Consistent with the results from HEK293-Langerin^+^ cells, mannosylated HJs significantly enhanced LC targeting compared to the non-glycosylated HJ (Figure 4D). By increasing the ligand quantity per scaffold from HJ-1xMan – HJ-3xMan and from HJ-1xTriMan – HJ-3xTriMan, LC targeting was enhanced by 2.8- and 1.9-fold, respectively. Additionally, HJ-3xMan significantly enhanced targeting compared to HJ-1xTriMan. Contrary to the binding studies with HEK293-Langerin^+^ cells, LCs were equally targeted by HJ-3xMan, HJ-2xTriMan and HJ-3xTriMan scaffolds after overnight incubation. When examining keratinocytes in the same suspension, the fluorescence signal for all HJ scaffolds was non-significant compared to untreated cells, suggesting a lack of binding.

Collectively, these results highlight the potential of mannosylated HJ scaffolds for targeting specifically LCs within the human epidermis.

### Topical delivery of mannosylated HJs to LCs within human skin

The drug delivery potential of mannosylated HJs to epidermal LCs was characterized by topical administration on human skin explants.

The outer layer of the human skin explants was perforated using a Dermaroller^TM^ and ATTO565-labeled non-glycosylated HJ, HJ-3xMan and HJ-3xTriMan were topically applied in PBS and incubated at 37 °C for 24 h. Afterwards, the skin was fixed in 4% formaldehyde, paraffin-embedded, and the FFPE-sections were stained for Langerin to determine HJ colocalization with LCs. By examining the epidermal region, LCs were dispersed between keratinocytes in the suprabasal layers and only marginally targeted by non-glycosylated HJ (Figure 5A). Upon delivery of HJ-3xMan and HJ-3xTriMan, both scaffolds colocalized with epidermal LCs without apparent keratinocyte interactions. Topical delivery of mannosylated HJs was conducted in skin explants from three different donors, and quantified by the percentage of colocalized HJ signal with the epidermal LCs using pixel-based colocalization (Figure 5B). Based on the mean colocalization from three donors, only 1.9% of non-glycosylated HJ colocalized with LCs, comparable to PBS treated skin. HJ-3xMan and HJ-3xTriMan both provided a significantly enhanced LC targeting of 11.6% and 17.6% colocalized signal, respectively. However, while our results indicate greater colocalization of HJ-3xTriMan with LCs compared to HJ-3xMan, this difference was found to be donor-dependent.

**Figure 5.**
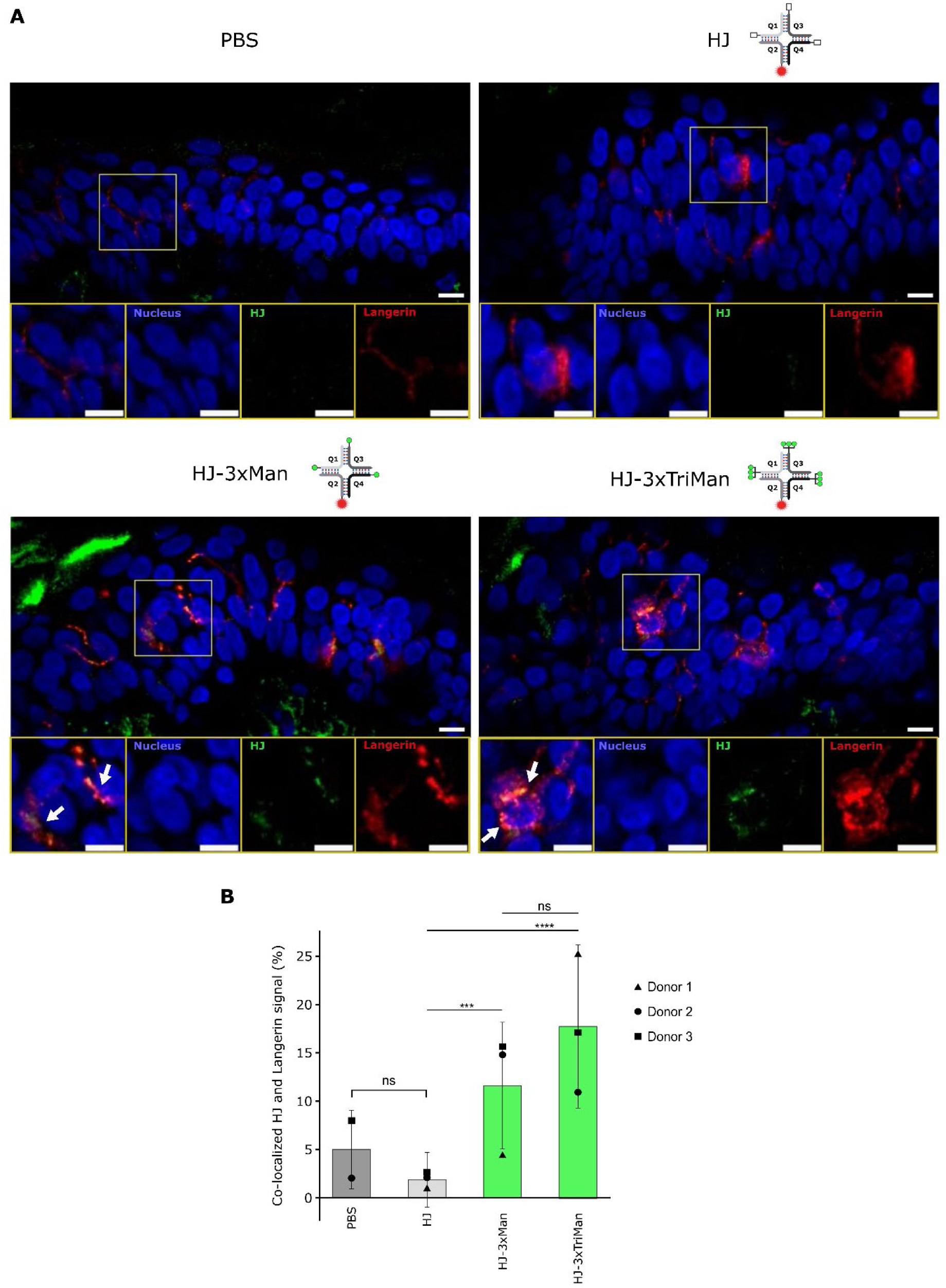
Topical delivery of mannosylated HJs to Dermarolled^TM^ human skin explants for targeting epidermal LCs. **A**) Representative microscopy images of epidermal skin after topical application of ATTO565-labeled, mannosylated HJ scaffolds (green): non-glycosylated HJ, HJ-3xMan and HJ-3xTriMan. PBS-treated skin was included as a negative control. The tissue was prepared as FFPEs and stained for LCs with anti-Langerin and secondary AF647-IgG antibody (red), and nucleus stained using DAPI (blue). A region of interest (ROI) was magnified (yellow square) and separated into individual fluorescent channels. Arrows indicate colocalizations between HJ and Langerin. Scale bar: 10 µm. **B**) Bar graphs showing the pixel-based colocalization of the fluorescent signal from the HJ scaffolds and the LCs was quantified in percentage (%) from three human skin donors (▲, ●, ■). Results expressed by the mean from n = 2 – 3 field-of-views (FOVs) of the epidermis from each donor and the error-propagated SD. Statistical analysis was conducted by a two-way T-test. P-value * = <0.05, ** = <0.01, *** = <0.001, ****= <0.0001.

Hereby we show the application of topically administered mannosylated HJ scaffolds for targeting LCs within the epidermal skin compartment.

### Effective delivery of antigenic peptides to Langerhans-like cells for T cell activation

Having established the effective targeting of HEK293-Langerin^+^ cells and ex vivo-derived human LCs, we proceeded to investigate tumor antigen delivery for facilitating T cell priming. For this purpose, an antigenic melanoma peptide gp100_154-162_ was conjugated on the mannosylated HJs.

Peptide conjugation onto the HJ oligonucleotides was achieved by SPAAC reaction between the azide-functionalized peptide and the DBCO-functionalized Q2 and Q4 strands (Figure 6A). The gp100 peptide was connected to the oligonucleotide strand using a lysosomally cleavable linker, making it susceptible to cathepsin B-mediated degradation intracellularly, leaving behind a *p*-amidobenzyloxycarbonyl (PAB) group, capable of undergoing spontaneous elimination to release free peptide^42^. The conjugate was purified by RP-HPLC and validated by denaturing PAGE (Supplementary Figure S7A-B). Efficient cleavage of the gp100 peptide from Q2 was confirmed by incubating the conjugate with cathepsin B for up to 5 h at 37 °C. Analysis by denaturing PAGE showed the oligonucleotide migrated faster after 0.5 h of cathepsin B incubation compared to the uncleaved sample at t = 0 h, indicating the removal of gp100 from Q2 (Supplementary Figure S7C).

**Figure 6.**
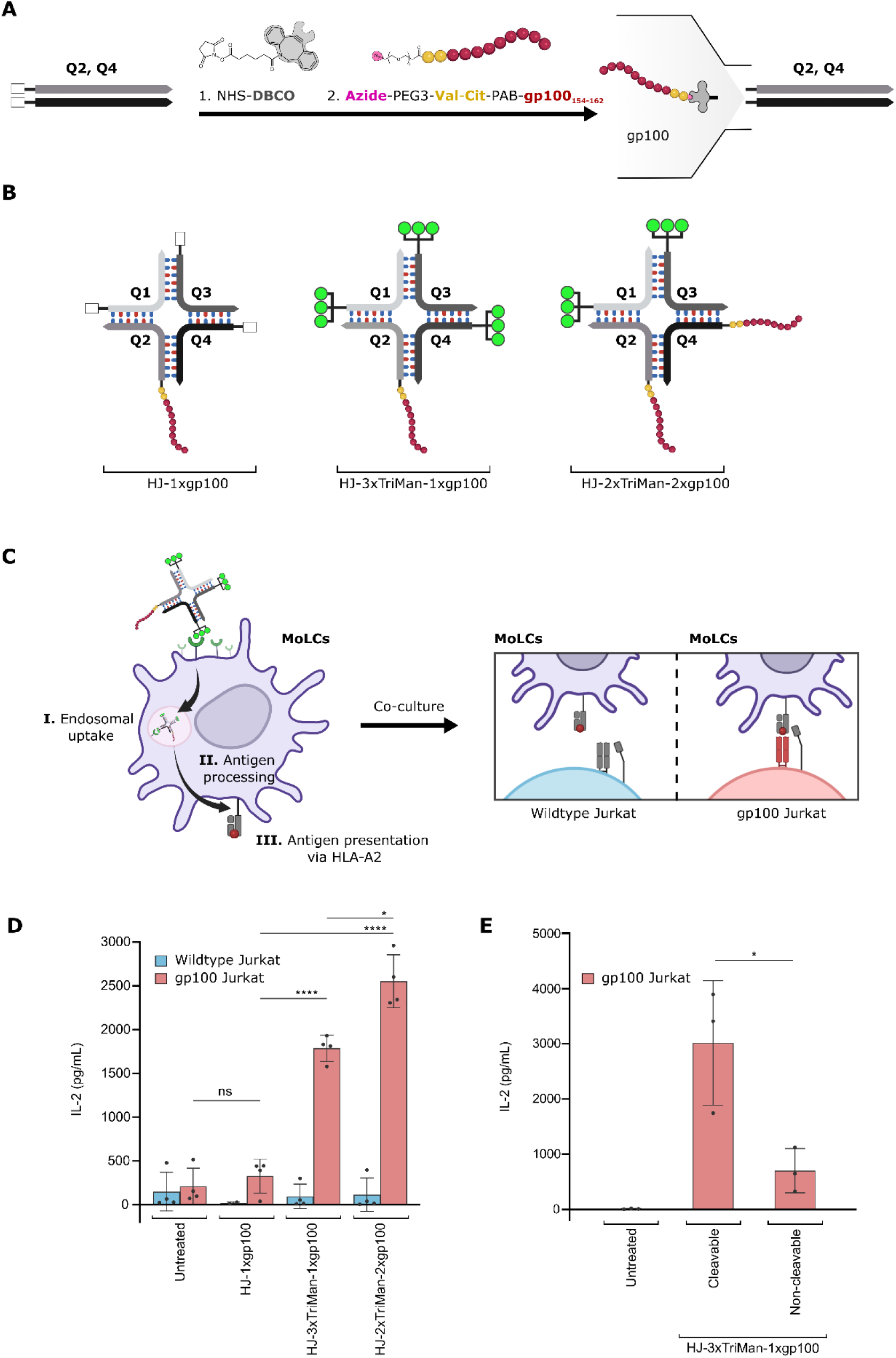
Delivery of gp100 on mannosylated HJ scaffolds into moLCs for antigen presentation. **A**) Reaction scheme for gp100-functionalization of Q2 and Q4 with NHS-DBCO (**1**), followed by SPAAC reaction to azide-modified (*pink*) Val-Cit-PAB-linked (*yellow)* gp100 (*red*) (**2**). **B**) Schematic representation of gp100-conjugated HJs employed for antigen presentation. **C**) Visual representation of the antigen-presentation assay using moLCs incubated with gp100-conjugated HJs to facilitate receptor-mediated endocytosis (**I**), followed by release of gp100 in the lysosomes (**II**) and presentation of gp100 on HLA-A2 (**III**). Subsequently, moLCs were co-cultured with wildtype- or gp100 Jurkat cells to verify antigen presentation by T cell activation. **D**) IL-2 concentration (pg/mL) in the co-culture media by ELISA. Results shown from four independent human donors were expressed by the mean ± SD. Statistical analysis was conducted by a one-way ANOVA followed by a Tukey’s post hoc test. **E**) Antigen-presentation assay comparing HJ-3xTriMan-1xgp100 assembled with either a cleavable linker (*Val-Cit*) or a non-cleavable linker. Results shown from three independent human donors were expressed by the mean ± SD. Statistical analysis was conducted by a two-way t test. p-value * = <0.05, ** = <0.01, *** = <0.001, ****= <0.0001.

The gp100-functionalized strands were assembled with TriMan ligands to form HJ displaying either one or two peptides per scaffold (HJ-3xTriMan-1xgp100 and HJ-2xTriMan-2xgp100, respectively) (Figure 6B). A non-glycosylated HJ containing one gp100 peptide (HJ-1xgp100) was included as a negative control.

Antigen presentation assays with gp100 were conducted using Langerin-expressing, monocyte-derived LCs (moLCs), differentiated from human monocytes as previously described^43,44^, using HLA-A2 positive donors. HJ scaffolds were efficiently internalized into moLCs, as determined in uptake studies using an EDTA-supplemented wash buffer to remove membrane-bound scaffolds, demonstrating comparable uptake of HJ-2xTriMan and HJ-3xTriMan (Supplementary Figure S8A-C).

Antigen loading of moLCs was achieved by incubating overnight with 2 µM of gp100-conjugated HJs in the presence of polyI:C to activate the cells. The following day, moLCs were co-cultured with wildtype- or gp100-specific T cell receptor expressing Jurkat cells (gp100 Jurkat) in a 1:1 ratio for 24 h (Figure 6C). T cell activation was quantified by interleukin-2 (IL-2) secretion, as determined by ELISA. To validate antigen-specific activation of Jurkat cells, unconjugated gp100 peptide was loaded onto HLA-A2 molecules of moLCs^45^ for 1 h at 37 °C which enhanced the IL-2 secretion during co-culture with gp100 Jurkat cells (Supplementary Figure S9).

The cleavable linker connecting gp100 peptide to the HJ scaffold was stable in the cell medium during the incubation, preventing the release of free antigenic peptides into the media (Supplementary Figure S10).

Upon moLC incubation with HJ-1xgp100, the IL-2 secretion was not significantly enhanced compared to untreated cells, indicating the scaffold itself does not facilitate antigen presentation to gp100 Jurkat cells (Figure 6D). In contrast, HJ-3xTriMan-1xgp100 and HJ-2xTriMan-2xgp100 loaded moLCs significantly enhanced IL-2 secretion during co-culture with gp100 Jurkat cells, compared to untreated moLCs and HJ-1xgp100. Additionally, doubling the peptide amount per scaffold using HJ-2xTriMan-2xgp100 enhanced IL-2 secretion by 1.4-fold compared to HJ-3xTriMan-1xgp100. Interestingly, both HJ-3xTriMan-1xgp100 and HJ-2xTriMan-2xgp100 were found to enhance the IL-2 secretion from gp100 Jurkat cells in lower concentration of 5 – 50 nM during moLC incubation (Supplementary Figure S11).

Next, we examined the effect of using a cleavable linker for delivery of gp100. To assess this, the cathepsin B-cleavable linker was compared to a non-cleavable linker using HJ-3xTriMan-1xgp100 in the antigen presentation assay (Figure 6E). Indeed, our findings showed the cleavable linker was necessary for efficient antigen presentation by moLCs to the gp100 Jurkat cells as determined by a 3-fold enhanced IL-2 secretion compared to the non-cleavable linker.

Our results demonstrate the ability to deliver gp100 to Langerhans-like cells with mannosylated HJ scaffolds, leading to effective T cell activation, while underlining the importance of cleavable linkers for efficient antigenic peptide release in lysosomes.

## Discussion

Harnessing DCs in situ to present antigens directly to immune cells has been proposed as an approach for immunization^8,9^. Taking advantage of our largest organ, the skin, we aimed to deliver a novel vaccine platform to epidermal LCs using a self-assembled nucleic acid-based scaffold. For this, we chose a serum-stable and non-immunogenic HJ scaffold, previously employed for therapeutic and diagnostic purposes^38–40^.

We identified the optimal carbohydrate and ligand configuration for targeting Langerin by screening five monosaccharides conjugated to each HJ strand, then assembling them in a mono-, bi-, and trivalent ligand arrangement. In vitro screening with HEK293-Langerin^+^ cells revealed significant binding preference towards Fuc, GlcNAc, and particularly Man, consistent with previous findings^19,46,47^. Monosaccharides significantly enhanced Langerin binding when assembled in bi- or trivalent arrangement on the HJ, corresponding with the glycocluster effect^5^, with HJ-3xMan showing the highest binding. To enhance Langerin binding, mannosylation was increased for each HJ strand using a TriMan ligand. Binding studies on HEK293-Langerin^+^ cells demonstrated an overall improvement with TriMan-conjugated HJ scaffolds, compared to monosaccharide Man, expectedly due to increased local ligand concentration. Interestingly, HJ-3xMan provided greater binding than HJ-1xTriMan, suggesting that optimal spacing of ligands on the HJ scaffold may enhance Langerin binding. This was also observed when targeting LCs from human epidermal cell suspensions. According to structural data, the Langerin protein forms trimers with an intradomain distance of 41.5 Å between each CRD^19^. This was substantiated in binding studies using molecular rulers made from DNA:PNA duplexes, demonstrating that Langerin ligands positioned 42 Å apart elicited the strongest binding, presumably facilitating chelation between two CRDs^48^. Considering the HJ scaffold spans approximately 40 Å for each 12 nt strand, ligand arrangement in bi- (Q1+Q3) or trivalent (Q1+Q3+Q4) display may bind multiple CRDs of the trimeric Langerin receptor, facilitating stronger binding. Taken together, our results suggest that in addition to using multivalent carbohydrate arrangements for enhanced Langerin binding, the HJ scaffold size may also offer structural advantages for targeting the receptor. However, further investigation is needed, as estimating ligand distance on the HJ is challenging due to the inherent scaffold and ligand linker flexibility^38^.

To evaluate primary LC targeting, epidermal cell suspensions were isolated from human skin and incubated with mannosylated HJs. The results revealed a valency-defined enhancement in LC targeting from mannosylated HJs without interaction with keratinocytes, yielding comparable targeting between HJ-3xMan, HJ-2xTriMan and HJ-3xTriMan. Additionally, mannosylated HJs were delivered to epidermal LCs following topical administration on Dermarolled^TM^ human skin explants. Quantitative colocalization of the HJ signal with LCs indicated enhanced targeting of both HJ-3xMan and HJ-3xTriMan, compared to non-glycosylated HJ. However, due to donor variations, no significant difference in LC targeting between HJ-3xMan and HJ-3xTriMan was observed, suggesting both mannosylation patterns are highly effective. The efficient LC targeting is likely due to the immediate exposure to the epidermis where they reside. Besides Dermarolling^TM^ for skin delivery, microneedle patches or intradermal injections may also be employed. However, these methods may release therapeutics directly into the dermis^49^, potentially redirecting mannosylated scaffolds towards other CLRs such as DC-SIGN or MR^7,8^. Alternatively, near-infrared lasers at low-energy settings can break the stratum corneum with minimal cell death around the laser pores, allowing antibody delivery to epidermal LCs^50^. Topical administration of DNA aptamers in cream-based formulations has recently been used to reduce inflammation by blocking CCL22 in murine skin^51^. However, murine skin thickness differs from human skin, and further research is needed to determine whether similar delivery vehicles are effective on human skin.

Our successful targeting of LCs prompted investigation into whether antigen delivery could be facilitated using mannosylated HJ. Therefore, we evaluated the delivery of a melanoma-associated antigenic peptide, gp100, conjugated to mannosylated HJs via a cleavable linker into moLCs. These cells were differentiated from human blood monocytes and co-cultured with gp100-recognizing T cells in an antigen-presentation assay. In this assay, moLCs were used to facilitate HLA-A2-specific complexation of gp100_154-162_^45^, instead of primary LCs, as HLA-A2-positive blood samples were more readily accessible, allowing for more biological replicates. Our findings showed that mannosylated HJs with one (HJ-3xTriMan-1xgp100) or two antigenic peptides (HJ-2xTriMan-2xgp100) were effectively delivered and presented by moLCs to activate T cells. Furthermore, HJ-2xTriMan-2xgp100 produced higher T cell activation than HJ-3xTriMan-1xgp100, likely attributed to a comparable uptake in moLCs but resulting in increased gp100 peptide presentation. However, antigen internalization and trafficking in moLCs may not be exclusive to Langerin-mediated uptake, as the expression of Man-binding DC-SIGN receptors has been reported, unlike in skin-derived LCs^43^. Since both Langerin and DC-SIGN can facilitate antigen presentation^24^, our results demonstrate potent antigen loading from mannosylated HJs following CLR-mediated uptake.

The application of glycosylated scaffolds for delivering tumor antigens to LCs has previously been accomplished using Lewis^Y^ glycan-modified peptides, resulting in successful antigen-presentation to CD8^+^ T cells^22^. However, cross-presentation of gp100 was not observed using a mannosylated peptide^23^. Conversely, our results show that mannosylated HJs effectively deliver gp100 into moLCs. The incorporation of a cleavable linker conjugated to the gp100 peptide on the HJ substantially improved IL-2 secretion compared to a non-cleavable linker. Protease-cleaved linkers were previously shown to improve antigen delivery to CD8^+^ T cells using viral-like particles coated with gp100, indicating the importance of intracellular release of free antigens for presentation^42^. Similarly, our findings suggest that an intracellularly cleaved, self-immolating linker may enhance lysosomal release of gp100, compared to unpredictably digested antigens from a non-cleavable linker, thereby improving antigen presentation to T cells. These linkers have not been thoroughly investigated for antigen delivery, as their initial development was focused on creating safer antibody-drug conjugates^52^.

In conclusion, we have demonstrated the modular nature of the HJ scaffold, highlighting its potential for carbohydrate-conjugated therapies. The plug-and-play assembly of carbohydrates and tumor antigenic peptides may serve as an effective drug delivery vehicle for immunotherapy to epidermal LCs, attainable by topical administration. Compared to invasive injectable administration routes, topical delivery for epidermal immunotherapy may improve patient compliance and reduce adverse effects. Interestingly, the modular HJ scaffold offers an easily expandable platform for drug delivery to other skin-residing cells through different targeting ligands. Future research could explore these possibilities, paving the way for innovative and less invasive skin therapeutic strategies.

## Materials and Methods

### Oligonucleotide sequences

Q1: 5’- NH_2_C_6_-mC+CmG+TmCmCmT+GmA+GmCmC –3’

Q2: 5’- NH_2_C_6_-mCmA+CmA+GmTmG+GmA+CmGmG –3’

Q3: 5’- NH_2_C_6_-mG+GmC+TmCmAmCmC+GmA+TmC –3’

Q4: 5’- NH_2_C_6_-mGmA+TmC+GmGmAmC+TmG+TmG –3’

Oligonucleotides strands (Q1-Q4) were synthesized with a 5’ amino modification with a six-carbon (C6) spacer arm. LNA and 2’OMe modified nucleotides are indicated with “+” and “m” prefixes, respectively. Oligonucleotides were purchased from Integrated DNA Technologies (IDT).

### Assembly of the HJ scaffold

Equimolar amounts of oligonucleotide strands (Q1-Q4) were mixed in a buffered solution (either in 200 mM potassium acetate (Sigma-Aldrich) or 1x phosphate buffered solution (PBS, Gibco)) and self-assembled under thermal annealing in a 90-min linear ramp from 70 °C to 4 °C.

### Gel electrophoresis

Assembled HJ scaffolds were visualized on native PAGE (8-16%) using AccuGel 29:1 (40%) (NationalDiagnostics) to the designated gel percentage in a total volume of 40 mL in 0.5x Tris-Borate-EDTA buffer (TBE, Gibco), catalyzed for polymerization with 320 µL ammonium persulfate (APS, Sigma-Aldrich) and 16 µL tetramethylethylenediamine (TEMED, Invitrogen^TM^).

Reaction products of conjugated oligonucleotide strands were visualized on denaturing PAGE (8-16%) using a mixture of UreaGel Diluent (National Diagnostics) and UreaGel Concentrate (National Diagnostics) to the designated gel percentage in a total volume of 40 mL in 1x TBE, catalyzed for polymerization with 320 µL APS and 16 µL TEMED. Gels were pre-run at 500V for 30 min prior to sample loading. For size comparison, an ultra low range DNA ladder (10-300 bp, Invitrogen^TM^) was included. The gels were stained in SYBR Gold (Invitrogen^TM^) for 15 min and scanned for fluorescence on the Amersham Typhoon biomolecular imager (Cytiva).

### Oligonucleotide Bioconjugation Chemistry

5’ amino modified oligonucleotides (Q1, Q3 and Q4) were DBCO-functionalized by reacting to a 10-fold molar excess of NHS-sulfo-DBCO (Jena Bioscience) in a buffered solution of 0.1 M HEPES (pH 8.3, Sigma-Aldrich) and 30 % DMSO (Sigma-Aldrich), shaking at RT overnight. DBCO-functionalized oligonucleotides were reacted with a 50-fold molar excess of azide-modified monosaccharides (2-azidoethyl *⍺⍺*-D-Galactoside (Gal), 2-azidoethyl N-acetyl-*⍺⍺*-D-Galactosamine (GalNAc), 2-azidoethyl *⍺⍺*-L-Fucopyranoside (Fuc), 2-azidoethyl 2-acetamido-2-deoxy-*⍺⍺*-D-Glucoside (GlcNAc), 2-azidoethyl *⍺⍺*-Mannoside-2-azido (Man), CarboSynUSA) in 0.1 M HEPES (pH 7.4) and 30 % DMSO, shaking at RT overnight.

Synthesis of the TriMan ligand was initiated by conjugating the 5’amino modified oligonucleotides (Q1, Q3, Q4) with a 10-fold molar excess of trivalent alkyne linker (NHS-PEG5-Tris-PEG4-alkyne, ConjuProbe) in a buffered solution of 0.1 M HEPES (pH 8.3) and 30% DMSO, shaking at RT overnight. Subsequently, the trialkyne-functionalized oligonucleotides were reacted to a 50-fold molar excess of azide-modified Man in a buffered solution of 0.1 M HEPES (pH 7.4) containing 440 μM copper(II)-sulfate (Sigma-Aldrich), 880 μM tris(benzyltriazolylmethyl)amine (TBTA, Cayman chemical), 16.6 μM ascorbic acid (Sigma-Aldrich) and 30-40% DMSO.

Fluorescent labeling of the 5’ amino modified Q2 strand was achieved by reaction to a 20-fold molar excess of NHS-functionalized fluorescent dyes (Cy5-NHS (Lumiprobe), ATTO647N-NHS (ATTO-TEC), ATTO565-NHS (ATTO-TEC)) in a buffered solution of 0.1 M HEPES (pH 8.3) containing 30-40% DMSO, shaking in the dark at RT overnight.

Antigenic peptide gp100_154-162_ (sequence: KTWGQYWQV, specificity: HLA-A*02:01, GENAXXON bioscience) was conjugated to the oligonucleotide strands in a 2-step one-pot reaction. First, a 4-fold molar excess of gp100 was reacted to the p-nitrophenyl ester (PNP) moiety of the cleavable crosslinker (azido-PEG3-Val-Cit-PAB-PNP, BroadPharm) in DMSO with 1% N,N-diisopropylethylamine (DIPEA, Sigma-Aldrich), shaking at RT for 3-4 h. Secondly, the reaction was spiked with DBCO-functionalized Q2 or Q4 in a 2-fold molar excess of the cleavable crosslinker in a buffered solution of 0.1 M HEPES (pH 7.4) containing 50% DMSO and incubated an additional 2-3 h.

The gp100_154-162_ peptide was correspondingly conjugated to the Q2 strand using a non-cleavable linker (azidobutyric acid NHS ester, Lumiprobe). Initially, 4-fold molar excess of gp100 was reacted with the non-cleavable linker a buffered solution of 0.1 M HEPES (pH 8.3) containing 50% DMSO, shaking at RT for 3-4 h. Afterwards, 2-fold molar excess of DBCO-functionalized Q2 was spiked into the reaction and incubated at RT an additional 2-3 h.

### Oligonucleotide purification by RP-HPLC

DBCO-, monosaccharide-, alkyne-, TriMan- and fluorophore-functionalized oligonucleotides were after each reaction subjected to ethanol precipitation and resuspended in nuclease-free water (Invitrogen^TM^) before RP-HPLC purification on an Agilent 1260 Infinity II series. Analytes were purified on a Kinetex EVO C18 column (150×4.6 mm, 100 Å, Phenomenex) with mobile phases A) 50 mM triethylammonium acetate (TEAA, PanReac AppliChem), 5% acetonitrile (MeCN, HPLC-grade, VWR chemicals) and B) 50 mM TEAA, 95% MeCN with a 0.4 mL/min flow rate. For chromatographic separation, a gradient elution was used with the mobile phase B composition: 0% from 0 to 2 min, 0 – 95% from 2 to 34 min, 95% from 34 to 37 min, 95 – 0% B from 37 – 40 min, 0 % B from 40 – 50 min. Reaction products were detected at 260 nm and fractions were collected, lyophilized and reconstituted in nuclease-free water.

### Cell culture

#### Langerin-expressing HEK293 cells

HEK293-Langerin^+^ cells were provided by Steffen Thiel, Department of Biomedicine, Aarhus University, Denmark. The cells were generated by transfecting HEK293F cells (Freestyle 293-F, Gibco) with expression vector pcDNA3.1/Zeo+ (GenScript) containing full-length open reading frames for Langerin (accession number NM_015717.5). The HEK293-Langerin^−^ cells were transfected with an empty expression vehicle. Cells were cultured in Freestyle^TM^ 293 expression medium (Gibco) supplemented with 10 % heat-inactivated FBS (Gibco), 1% Penicillin-Streptomycin (Gibco) and 500 *μμ*g/mL Zeocin (Invitrogen^TM^), cultured at 37 °C. The adherent cells were detached from culture flasks using 0.05% Trypsin-EDTA (Gibco). Langerin expression was validated by staining with anti-Langerin antibody (APC-conjugated, clone 10E2, Biolegend, cat # 352206).

#### Human epidermal cell suspensions

Healthy human skin samples from breast and abdomen reduction surgeries were obtained from the Department for Plastic, Reconstructive and Aesthetic Surgery of the Medical University Innsbruck, Austria, after informed consent and approval by the local ethics committee (AN5003 323/4.10 403/5.10 (4470a)). Dermatomized skin samples (1 mm thickness) were incubated for 30 min at 4 °C in RPMI-1640 (PAN-Biotech) supplemented with 50 µg/mL gentamicin (Gibco) for disinfection. Skin samples were cut into 1×1 cm pieces and incubated floating on RPMI-1640 containing 1.5 IU/mL Dispase II (Roche), and 0.1 % Trypsin (Sigma-Aldrich) at 4 °C overnight. The epidermis was peeled off and cut into small pieces with scissors. To generate a single cell suspension, skin pieces were passed through a 100 µm cell strainer (Corning) followed by a 40 µm cell strainer to remove cell clumps. The epidermal cells were cultured at 37 °C in R10 medium (RPMI1640 (PAN-Biotech), 2 mM L-glutamine (PAN-Biotech), 50 µg/mL gentamycin (Gibco), heat-inactivated FBS (PAN-Biotech)) supplemented with 200 IU/mL recombinant human granulocyte-macrophage colony stimulating factor (GM-CSF, specific activity of approx. 5.6 × 10^6^ IU/mg, LEUKINE®, Sanofi), to keep LCs viable.

#### Generation of moLCs

Peripheral blood from healthy human donors was obtained from the local blood bank after informed consent and approval by the local ethics committee (1265/2019). Blood was diluted 1:1 in 1x PBS, layered onto Lymphoprep (1.077 g/mL; STEMCELL Technologies) to isolate peripheral blood mononuclear cells by density gradient centrifugation at 800 g for 30 min at RT. CD14^+^ monocytes were purified by positive magnetic cell separation using anti-CD14 magnetic particles (clone MφP9, BD Biosciences) according to manufacturer’s instructions. Purity (>90 %) was confirmed by flow cytometry on CytoFLEX S (Beckman Coulter Life Sciences) using an anti-CD14 antibody (APC-conjugated, clone HCD14, Biolegend, cat # 325608).

Monocytes were differentiated to moLC as previously described by co-culture on a monolayer of OP9 cells expressing delta like canonical Notch ligand 4 (OP9-DLL4, kindly provided by Juan Carlos Zúñiga-Pflücker, Sunnybrook Research Institute, Department of Immunology, University of Toronto, Canada)^44^. The OP9-DLL4 feeder cells were cultured in R10 medium and assessed for DLL4 expression by flow cytometry using an anti-DLL4 antibody (APC-conjugated, clone HMD4-1, Biolegend, cat #130814). For the co-culture, a 12-well plate was seeded with 6 × 10^4^ OP9-DLL4 cells/well. After 24 h, 2.4 × 10^5^ CD14^+^ monocytes were added to each well with 280 IU/mL recombinant human GM-CSF and 10 ng/mL recombinant human TGF-β1 (PeproTech) and incubated at 37 °C. After 72 h, the differentiated moLCs were harvested by rinsing the wells with cold sterile 1x PBS.

### Flow Cytometry

#### Binding and uptake in Langerin-expressing HEK293 cells

Binding assays on HEK293 cells were conducted with Cy5-labeled HJ scaffolds. 2.0 × 10^5^ cells/well were seeded into U-bottomed 96-well plates in Freestyle^TM^ supplemented medium, centrifuged (1000 rpm, 5 min) and resuspended in a cell staining buffer (20 mM HEPES (pH 7.4) supplemented with 5 mM CaCl_2_ (Sigma-Aldrich), 5 mM MgCl_2_ (Sigma-Aldrich), 150 mM NaCl (VWR chemicals) and 0.5% BSA (Sigma-Aldrich)). The suspensions were incubated with the HJs in the dark at RT for 30 min, washed thrice in the cell staining buffer and analyzed by flow cytometry.

Uptake kinetics in HEK293 cells were measured by incubating Cy5-labeled HJ scaffolds for 5 – 300 min in Freestyle^TM^ supplemented medium at 37 °C, washed thrice in 20 mM HEPES (pH 7.4) with 20 mM EDTA (Sigma-Aldrich). Cells were resuspended and analyzed in 20 mM HEPES (pH 7.4) on the Novocyte Acea flow cytometer (Agilent).

#### HJ targeting in epidermal cell suspension

For this purpose, 7.5 × 10^5^ human epidermal cells containing LCs were cultured in R10 medium supplemented with 200 IU/mL GM-CSF in a U-bottom 96-well plate. Then 20 nM of ATTO647N-labeled HJs were added to the cell suspensions and incubated overnight at 37 °C. Cells were centrifuged (1200 rpm, 4 °C, 5 min) and washed in WB buffer (1x PBS (Gibco), 1% BSA (SERVA), 5 mM EDTA (LONZA), 20 µg/mL DNase I (Sigma-Aldrich)) and analyzed on the CytoFLEX S flow cytometer (Beckman Coulter). Dead cells were excluded from analysis using SYTOX™ Orange dead cell stain (Thermo Fisher Scientific). Nonspecific FcR-mediated binding of antibodies was prevented by using human Fc blocking reagent (BD Biosciences). Staining with anti-CD45 (BV650-conjugated, clone HI30, Biolegend, cat # 304044), anti-CD1a (FITC-conjugated, clone HI149, BD Biosciences, cat # 555806), and anti-HLA-DR (PE-Cy7-conjugated, clone L243, Biolegend, cat # 307616) antibodies was performed at 4 °C for 15 min in 1x PBS supplemented with 1% BSA.

#### Uptake assays in moLCs

To assess if the WB buffer removes membrane-bound mannosylated HJ from the moLCs, cells were incubated at 4 °C with 100 nM ATTO647N-labeled HJ-3xTriMan for 22 h, then washed with either WB or Hanks’ balanced salt solution (HBSS) and analyzed by flow cytometry.

The uptake kinetics for HJ-2xTriMan and HJ-3xTriMan was subsequently determined by seeding 1.5 × 10^4^ moLCs into a U-bottom 96-well plate in R10 medium supplemented with 280 IU/mL GM-CSF, incubated with 25 nM ATTO647N-labeled HJ for 1 – 22 h at 37 °C. Afterwards, moLCs were washed in WB buffer and analyzed on the CytoFLEX S flow cytometer after staining with anti-CD45 (BV510-conjugated, clone HI30, Biolegend, cat # 30403), anti-Langerin (VioBlue-conjugated, clone MB22-9F5, Miltenyi Biotec) and anti-CD1a (FITC-conjugated, clone HI149, BD Biosciences, cat # 555806) antibodies at 4 °C for 15 minutes in 1x PBS supplemented with 1% BSA. Dead cells were excluded from analysis using SYTOX™ Orange dead cell stain (Thermo Fisher Scientific).

#### Flow cytometry data analysis

Data acquired from flow cytometry was analyzed either on the NovoExpress V.1.6.2 (Agilent) or FlowJo V.9 software (BD Biosciences).

### Microscopy of intracellular uptake

In 8-well chambers (Ibi-treated µ-slide, IBIDI GMBH), 5 × 10^4^ HEK293-Langerin^+^ cells were seeded in Freestyle 293^TM^ supplemented medium and incubated overnight at 37 °C. Cells were incubated with 200 nM Cy5-labeled HJs with Lysotracker^TM^ Green DND-26 (Invitrogen^TM^). After 24 h, cells were stained with Hoechst 33342 (Thermo Scientific) for 30 min, washed in medium and imaged on a confocal microscope (LSM700, Zeiss) with a 63x oil-objective. Images were adjusted for brightness and contrast equally in Fiji ImageJ software for improved visualization.

### Topical skin delivery of HJs

Human skin explants were obtained from healthy female patients undergoing breast, belly and back reduction surgery kindly provided by Claus Johansen from the Department of Dermatology and Venereology, Aarhus University Hospital, Aarhus, Denmark. The skin was washed with 1xPBS, trimmed for subcutaneous tissue and the stratum corneum was perforated using a 0.5 mm Dermaroller^TM^ (Argador). The skin was cut into square pieces of approximately 1.5 cm^2^ and placed in DMEM Glutamax (Gibco) supplemented with 1% penicillin-streptomycin (Gibco), 0.5% gentamicin (Gibco) and 10% FBS (Gibco). ATTO565-labeled HJ were topically applied in 1x PBS to the epidermal side of the skin and incubated for 24 h at 37 °C. After incubation, punch biopsies (4 mm) were extracted from the skin and dehydrated, fixed in 4% formaldehyde then embedded in paraffin wax. The tissue blocks were sectioned (approximately 10 µm thick) and the tissue slices were stained with mouse anti-Langerin antibody (clone 12D6, Cell Maque, cat 392M-15) and a secondary donkey anti-mouse AF647-IgG (Alexa Fluor 647-conjugateed, Invitrogen, cat A-315771), then nucleus stained with DAPI-supplemented mounting medium (Fluoroshield mounting medium, Abcam). The tissue samples were imaged on a confocal microscope (LSM700, Zeiss) using a 63x oil-objective. Images were taken from 10 z-slices, equally adjusted for brightness and contrast in Fiji ImageJ software. The 10 z-slices were Z projected to one image using maximum intensity. The Z projected images were hard thresholded in Fiji ImageJ to create binary images to display colocalization of HJ and Langerin signal. From the binary images, colocalization was determined by subtracting the binary images. A home-written python script was developed for automatically integrating colocalized areas, and calculating the percentage of targeting for each technical replicate *P_n_* from:

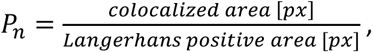

Where *colocalized area* [*px*] are the integrated colocalized areas and *Langerhans positive area* [*px*] is the integrated LCs positive area, based on the Langerin signal.

For each donor, n = 2-3 technical samples were integrated to obtain their respective mean and SD. Based on that, from three donors a collective mean and the error-propagated SD was calculated.

### Antigen presentation assay

Gp100-specific Jurkat cells were generated by lentiviral introduction of the pCDH_gp100TCR_IRES-GFP_puro vector kindly provided by David Schrama (Department of Dermatology, Venerology and Allergology, University Hospital Würzburg, Germany). Successful transduction of Jurkat cells was assured by antibiotic selection with puromycin and GFP-positivity. The gp100-specific T cell receptor expressed by the vector-carrying Jurkat cells is directed against the gp100_154-162_ peptide presented on HLA-A2. Recognition of the HLA-A2-restricted gp100 peptide by the T cell receptor induces IL-2 secretion of Jurkat cells.

Blood donors tested positive for HLA-A2 genotype were used for differentiation of moLCs. For the antigen-presentation assay, 3.0 × 10^4^ freshly harvested moLCs were seeded into a U-bottomed 96-well plate in R10 medium and incubated with HJs overnight at 37 °C, in the presence of 280 U/mL GM-CSF and 20 ng/µL poly I:C (Sigma-Aldrich), to keep the moLCs viable and induce activation. Afterwards moLCs were co-cultured with 3.0 × 10^4^ gp100-specific or wildtype Jurkat cells in R10 medium supplemented with 280 U/mL GM-CSF, incubated for 24 h at 37 °C. As a positive control, untreated moLCs were incubated with free gp100_154-162_ for 1 h at 37 °C prior to co-culturing with Jurkat cells. Co-culture supernatants were quantified for their IL-2 concentration using ELISA MAX™ Deluxe Set Human IL-2 (Biolegend) according to manufacturer’s instructions.

### Cathepsin B cleavage assay

25 mM cathepsin B (human liver, ≥ 1,500 units/mg protein, Sigma-Aldrich), 15 mM DTT (Thermo Scientific), 7.5 mM EDTA incubated for 15 min at RT, then diluted in 37 °C-preheated 30 mM sodium acetate (pH 5.0, Sigma-Aldrich) and spiked with 450 pmol gp100-conjugated Q2. The solution was incubated at 37 °C and aliquots were taken out after t = 0, 30, 60, 180, 240 and 300 min, then placed on dry ice. The aliquots were visualized on an 8% denaturing PAGE to visualize the gp100 cleavage from Q2.

### Statistics and figures

Graphs were prepared and statistically analyzed with Graphpad Prism version 10.1.2. Figures were created using BioRender, Inkscape and PerkinElmer Chemdraw version 22.2.0.

## References

1. Zhang, L. et al. The therapeutic prospects of N-acetylgalactosamine-siRNA conjugates. Front. Pharmacol. 13, 1–13 (2022).

2. Debacker, A. J., Voutila, J., Catley, M., Blakey, D. & Habib, N. Delivery of Oligonucleotides to the Liver with GalNAc: From Research to Registered Therapeutic Drug. Mol. Ther. 28, 1759–1771 (2020).

3. Zelensky, A. N. & Gready, J. E. The C-type lectin-like domain superfamily. FEBS J. 272, 6179–6217 (2005).

4. Lee, Y. C. & Lee, R. T. Carbohydrate-Protein Interactions: Basis of Glycobiology. Acc. Chem. Res. 28, 321–327 (1995).

5. Lundquist, J. J. & Toone, E. J. The cluster glycoside effect. Chem. Rev. 102, 555–578 (2002).

6. Tjandra, K. C. & Thordarson, P. Multivalency in Drug Delivery–When Is It Too Much of a Good Thing? Bioconjug. Chem. 30, 503–514 (2019).

7. van Dinther, D. et al. Targeting C-type lectin receptors: a high-carbohydrate diet for dendritic cells to improve cancer vaccines. J. Leukoc. Biol. 102, 1017–1034 (2017).

8. Stoitzner, P., Romani, N., Rademacher, C., Probst, H. C. & Mahnke, K. Antigen targeting to dendritic cells: Still a place in future immunotherapy? Eur. J. Immunol. 52, 1909–1924 (2022).

9. Steinman, R. M. Decisions about dendritic cells: Past, present, and future. Annu. Rev. Immunol. 30, 1–22 (2012).

10. Engering, A. et al. The Dendritic Cell-Specific Adhesion Receptor DC-SIGN Internalizes Antigen for Presentation to T Cells. J. Immunol. 168, 2118–2126 (2002).

11. Merad, M., Ginhoux, F. & Collin, M. Origin, homeostasis and function of Langerhans cells and other langerin-expressing dendritic cells. Nat. Rev. Immunol. 8, 935–947 (2008).

12. Romani, N., Clausen, B. E. & Stoitzner, P. Langerhans cells and more: langerin-expressing dendritic cell subsets in the skin. Immunol. Rev. 234, 120–141 (2010).

13. Birbeck, M. S., Breathnach, A. S. & Everall, J. D. An Electron Microscope Study of Basal Melanocytes and High-Level Clear Cells (Langerhans Cells) in Vitiligo. J. Invest. Dermatol. 37, 51–64 (1961).

14. Valladeau, J. et al. Langerin, a Novel C-Type Lectin Specific to Langerhans Cells, Is an Endocytic Receptor that Induces the Formation of Birbeck Granules. Immunity 12, 71–81 (2000).

15. Idoyaga, J. et al. Cutting Edge: Langerin/CD207 Receptor on Dendritic Cells Mediates Efficient Antigen Presentation on MHC I and II Products In Vivo. J. Immunol. 180, 3647–3650 (2008).

16. Idoyaga, J. et al. Specialized role of migratory dendritic cells in peripheral tolerance induction. J. Clin. Invest. 123, 844–854 (2013).

17. Bouteau, A. et al. DC Subsets Regulate Humoral Immune Responses by Supporting the Differentiation of Distinct Tfh Cells. Front. Immunol. 10, 1–15 (2019).

18. Hanske, J. et al. Intradomain Allosteric Network Modulates Calcium Affinity of the C-Type Lectin Receptor Langerin. J. Am. Chem. Soc. 138, 12176–12186 (2016).

19. Feinberg, H., Powlesland, A. S., Taylor, M. E. & Weis, W. I. Trimeric Structure of Langerin. J. Biol. Chem. 285, 13285–13293 (2010).

20. Wamhoff, E.-C. et al. A Specific, Glycomimetic Langerin Ligand for Human Langerhans Cell Targeting. ACS Cent. Sci. 5, 808–820 (2019).

21. Schulze, J. et al. A Liposomal Platform for Delivery of a Protein Antigen to Langerin-Expressing Cells. Biochemistry 58, 2576–2580 (2019).

22. Fehres, C. M. et al. Langerin-mediated internalization of a modified peptide routes antigens to early endosomes and enhances cross-presentation by human Langerhans cells. Cell. Mol. Immunol. 14, 360–370 (2017).

23. Li, R.-J. E. et al. Targeting of the C-Type Lectin Receptor Langerin Using Bifunctional Mannosylated Antigens. Front. Cell Dev. Biol. 8, 1–8 (2020).

24. Fehres, C. M. et al. Cross-presentation through langerin and DC-SIGN targeting requires different formulations of glycan-modified antigens. J. Control. Release 203, 67–76 (2015).

25. Duinkerken, S. et al. Glyco-dendrimers as intradermal anti-tumor vaccine targeting multiple skin DC subsets. Theranostics 9, 5797–5809 (2019).

26. Neuhaus, K. et al. Asymmetrically Branched Precision Glycooligomers Targeting Langerin. Biomacromolecules 20, 4088–4095 (2019).

27. van Dongen, M. A., Dougherty, C. a & Banaszak Holl, M. M. Multivalent Polymers for Drug Delivery and Imaging: The Challenges of Conjugation. Biomacromolecules 15, 3215–3234 (2014).

28. Wang, L. et al. Therapeutic peptides: current applications and future directions. Signal Transduct. Target. Ther. 7, 48 (2022).

29. Jiang, S., Ge, Z., Mou, S., Yan, H. & Fan, C. Designer DNA nanostructures for therapeutics. Chem 7, 1156–1179 (2021).

30. Poppleton, E. et al. RNA origami: design, simulation and application. RNA Biol. 20, 510–524 (2023).

31. Kawasaki, A. M. et al. Uniformly modified 2’-deoxy-2’-fluoro-phosphorothioate oligonucleotides as nuclease-resistant antisense compounds with high affinity and specificity for RNA targets. J. Med. Chem. 36, 831–841 (1993).

32. Cummins, L. L. et al. Characterization of fully 2’-modified oligoribonucleotide hetero- and homoduplex hybridization and nuclease sensitivity. Nucleic Acids Res. 23, 2019–2024 (1995).

33. Elmén, J. et al. Locked nucleic acid (LNA) mediated improvements in siRNA stability and functionality. Nucleic Acids Res. 33, 439–447 (2005).

34. Koshkin, A. A. et al. LNA (Locked Nucleic Acids): Synthesis of the adenine, cytosine, guanine, 5-methylcytosine, thymine and uracil bicyclonucleoside monomers, oligomerisation, and unprecedented nucleic acid recognition. Tetrahedron 54, 3607–3630 (1998).

35. Kratschmer, C. & Levy, M. Effect of Chemical Modifications on Aptamer Stability in Serum. Nucleic Acid Ther. 27, 335–344 (2017).

36. Teodori, L., Omer, M. & Kjems, J. RNA nanostructures for targeted drug delivery and imaging. RNA Biol. 21, 1–19 (2024).

37. Yip, T., Qi, X., Yan, H. & Chang, Y. Therapeutic applications of RNA nanostructures. RSC Adv. 14, 28807–28821 (2024).

38. Andersen, V. L. et al. A self-assembled, modular nucleic acid-based nanoscaffold for multivalent theranostic medicine. Theranostics 9, 2662–2677 (2019).

39. Omer, M., Andersen, V. L., Nielsen, J. S., Wengel, J. & Kjems, J. Improved Cancer Targeting by Multimerizing Aptamers on Nanoscaffolds. Mol. Ther. - Nucleic Acids 22, 994–1003 (2020).

40. Teodori, L. et al. Plug-and-play nucleic acid-mediated multimerization of biparatopic nanobodies for molecular imaging. Mol. Ther. - Nucleic Acids 35, 102305 (2024).

41. Gerke, C. et al. Sequence-Controlled Glycopolymers via Step-Growth Polymerization of Precision Glycomacromolecules for Lectin Receptor Clustering. Biomacromolecules 18, 787–796 (2017).

42. Carl, P. L., Chakravarty, P. K. & Katzenellenbogen, J. A. A novel connector linkage applicable in prodrug design. J. Med. Chem. 24, 479–480 (1981).

43. Bellmann, L. et al. Notch-Mediated Generation of Monocyte-Derived Langerhans Cells: Phenotype and Function. J. Invest. Dermatol. 141, 84–94.e6 (2021).

44. Milne, P. et al. Hematopoietic origin of Langerhans cell histiocytosis and Erdheim-Chester disease in adults. Blood 130, 167–175 (2017).

45. Barker, A. B. H. et al. Identification of a novel peptide derived from the melanocyte-specific gp100 antigen as the dominant epitope recognized by an HLA-A2.1-restricted anti-melanoma CTL line. Int. J. Cancer 62, 97–102 (1995).

46. Stambach, N. S. & Taylor, M. E. Characterization of carbohydrate recognition by langerin, a C-type lectin of Langerhans cell. Glycobiology 13, 401–410 (2003).

47. Holla, A. & Skerra, A. Comparative analysis reveals selective recognition of glycans by the dendritic cell receptors DC-SIGN and Langerin. Protein Eng. Des. Sel. 24, 659–669 (2011).

48. Bachem, G., et al. Rational Design of a DNA-Scaffolded High-Affinity Binder for Langerin. Angew. Chemie Int. Ed. 59, 21016–21022 (2020).

49. Badran, M. M., Kuntsche, J. & Fahr, A. Skin penetration enhancement by a microneedle device (Dermaroller®) in vitro: Dependency on needle size and applied formulation. Eur. J. Pharm. Sci. 36, 511–523 (2009).

50. Tripp, C. H. et al. Laser-assisted epicutaneous immunization to target human skin dendritic cells. Exp. Dermatol. 30, 1279–1289 (2021).

51. Jonczyk, A. et al. Topical application of a CCL22-binding aptamer suppresses contact allergy. Mol. Ther. Nucleic Acid 35, 102254 (2024).

52. Bargh, J. D., Isidro-Llobet, A., Parker, J. S. & Spring, D. R. Cleavable linkers in antibody-drug conjugates. Chem. Soc. Rev. 48, 4361–4374 (2019).

