## Supplementary Figures for "Targeting Langerhans cells using a modular mannosylated nucleic acid-based vaccine platform"

### Supplementary Information

For manuscript titled: *Targeting Langerhans cells using a modular mannosylated nucleic acid-based vaccine platform.*

#### Table of Contents

Supplementary Figures


S1 ... 2
S2 ... 3
S3 ... 4
S4 ... 5
S5 ... 6
S6 ... 10
S7 ... 12
S8 ... 13
S9 ... 14
S10 ... 15
S11 ... 16

### Supplementary Figure S1

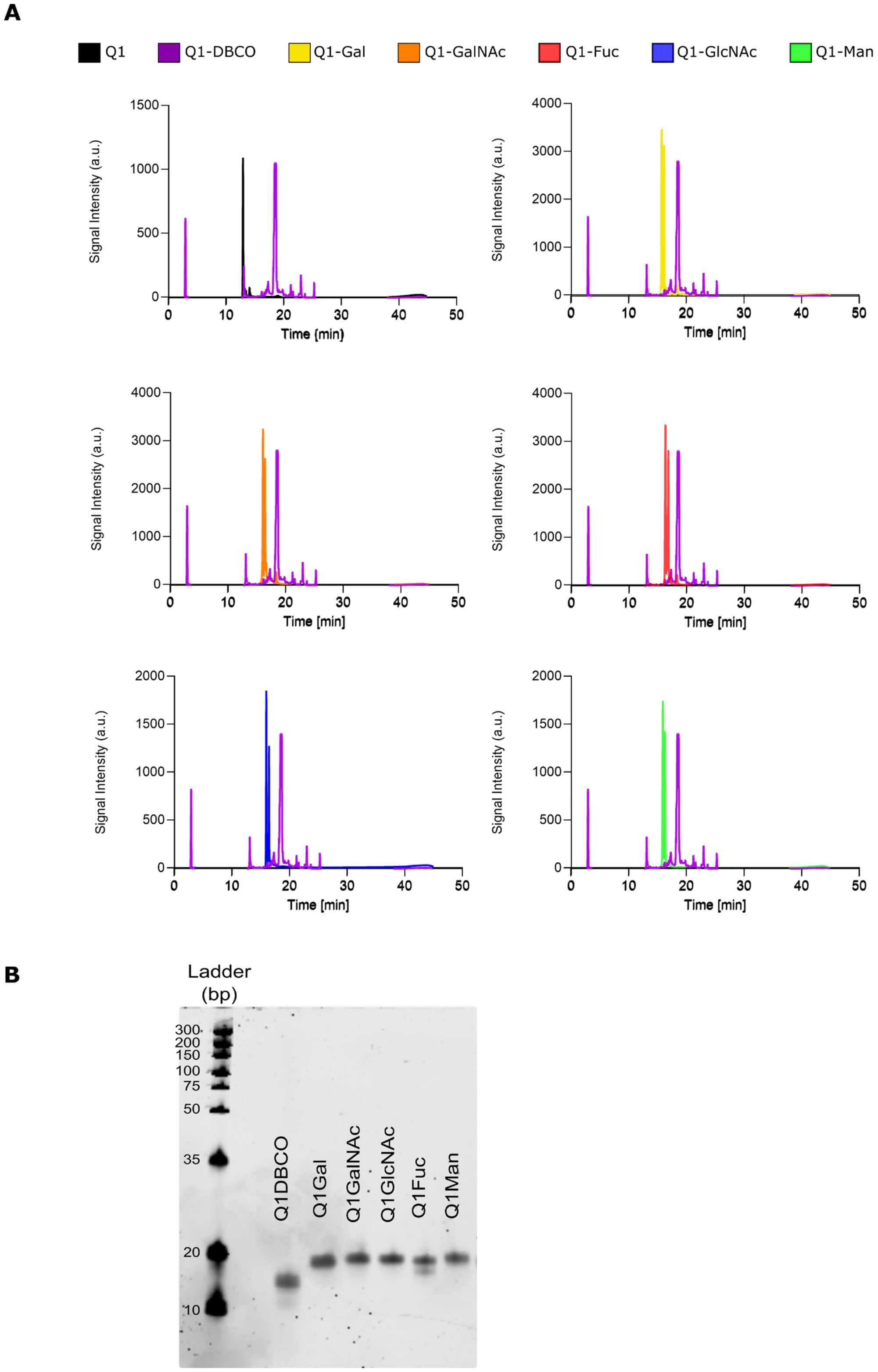


**Supplementary Figure S1. Purification and PAGE analysis of monosaccharide-conjugated Q1 strand. A)** RP-HPLC chromatograms (absorbance: 260 nm) for purification of monosaccharide-conjugated strands, exemplified by Q1 (Rt: 12.9 min) following DBCO-functionalization (Rt = 18.7 min). The Q1-DBCO conjugate was reacted with azide-functionalized monosaccharides (Gal, GalNAc, Fuc, GlcNAc, and Man) and subsequently purified by HPLC (*Rt* = 15.7, 16.1, 16.3, 16.0, and 16.2 min, respectively). **B)** HPLC-purified Q1 conjugates visualized on a denaturing 16% PAGE stained with SYBR Gold, showing the gel shift from Q1-DBCO to Q1-Gal, Q1-GalNAc, Q1 -GlcNAc, Q1-Fuc, and Q1-Man.

#### Supplementary Figure S2

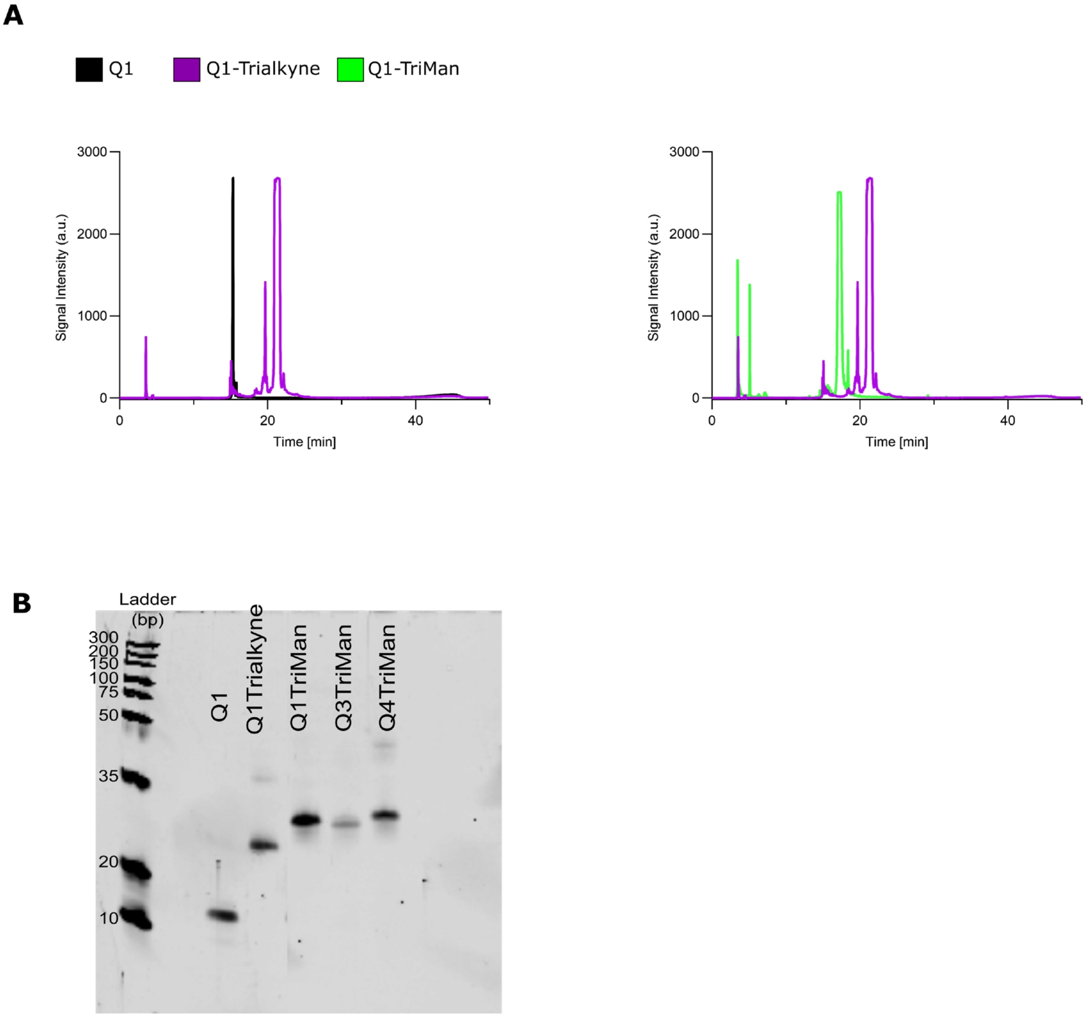


**Supplementary Figure S2. Purification and PAGE analysis of TriMan-conjugated oligonucleotide strands**. **A**) HPLC chromatograms (absorbance: 260 nm) for purification of TriMan-conjugated strands, exemplified by Q1 (Rt: 12.9 min) following trialkyne linker-functionalization (NHS-PEG5-Tris-PEG4-Alkyne$_3$, Rt: 21.5 min). The Q1-Trialkyne was reacted with an excess of azide-modified Man to yield Q1-TriMan (Rt: 17.1 min). **B**) HPLC-purified TriMan conjugates visualized in a denaturing 16% PAGE stained with SYBR Gold, showing the gel shift from Q1 and Q1-Trialkyne to Q1-TriMan, Q3-TriMan and Q4-TriMan.

#### Supplementary Figure S3

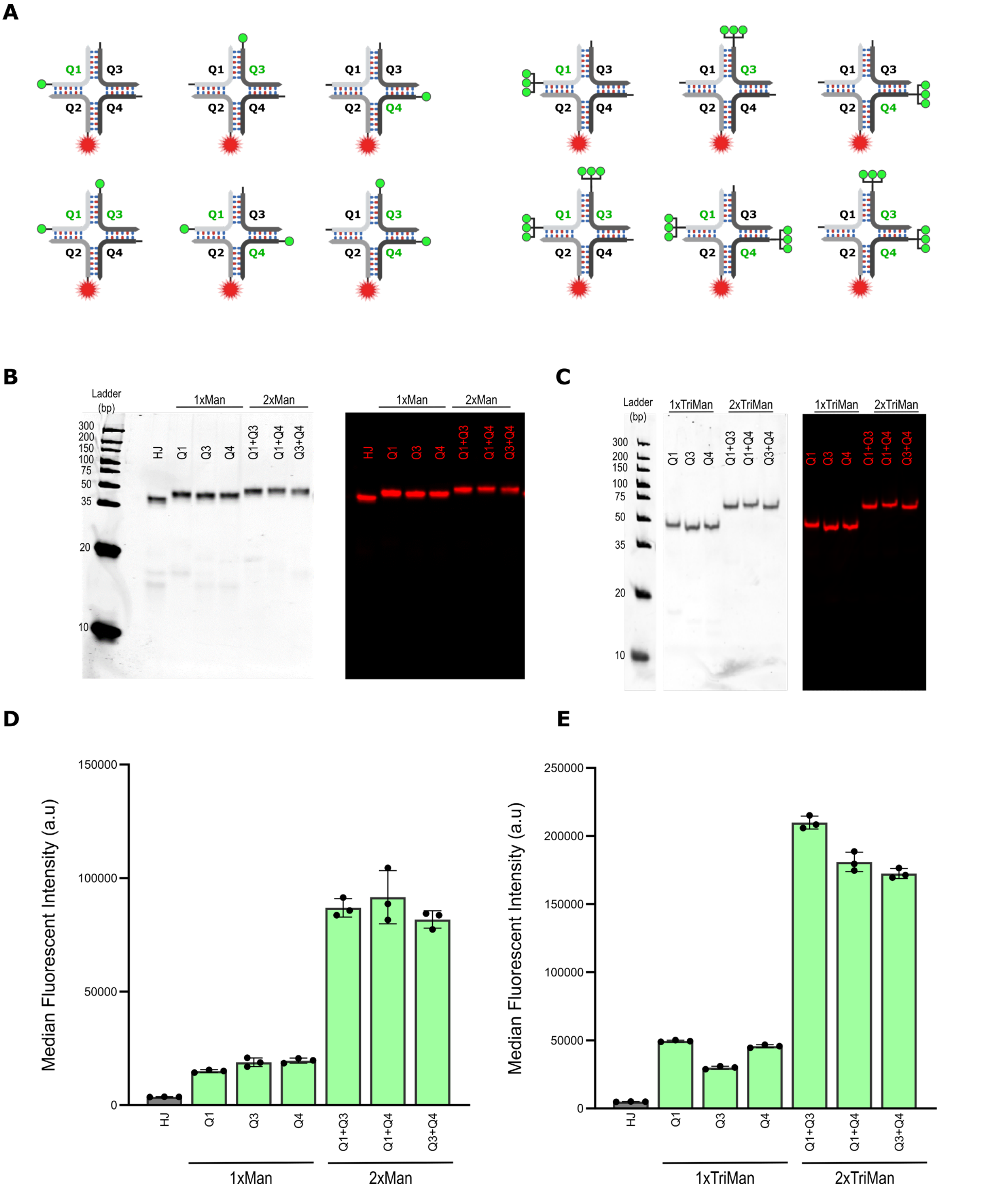


**Supplementary Figure S3. Evaluation of binding of mannosylated HJs on Langerin-expressing HEK293 Cells**. **A)** Schematic representation of different arrangements of Cy5-labeled mannosylated HJs. Man- and TriMan-conjugated oligonucleotides (green circles) were positioned on the HJ scaffold either monovalently on Q1, Q3, or Q4 or bivalently on Q1+Q3, Q1+Q4, or Q3+Q4, as denoted by green text color. **B-C)** Image of 16% native PAGE of Man- and TriMan-conjugated HJs stained with SYBR Gold and scanned for Cy5-fluorescence signal (left and right, respectively). **D-E)** Binding of Cy5-labeled mannosylated HJs to HEK293-Langerin⁺ cells at 50 nM measured by flow cytometry, with non-glycosylated HJ (grey) included as a control. Bar graphs of the MdFI displayed from the flow cytometry analysis shown from n = 3 technical replicates, expressed by the mean ± SD.

### Supplementary Figure S4

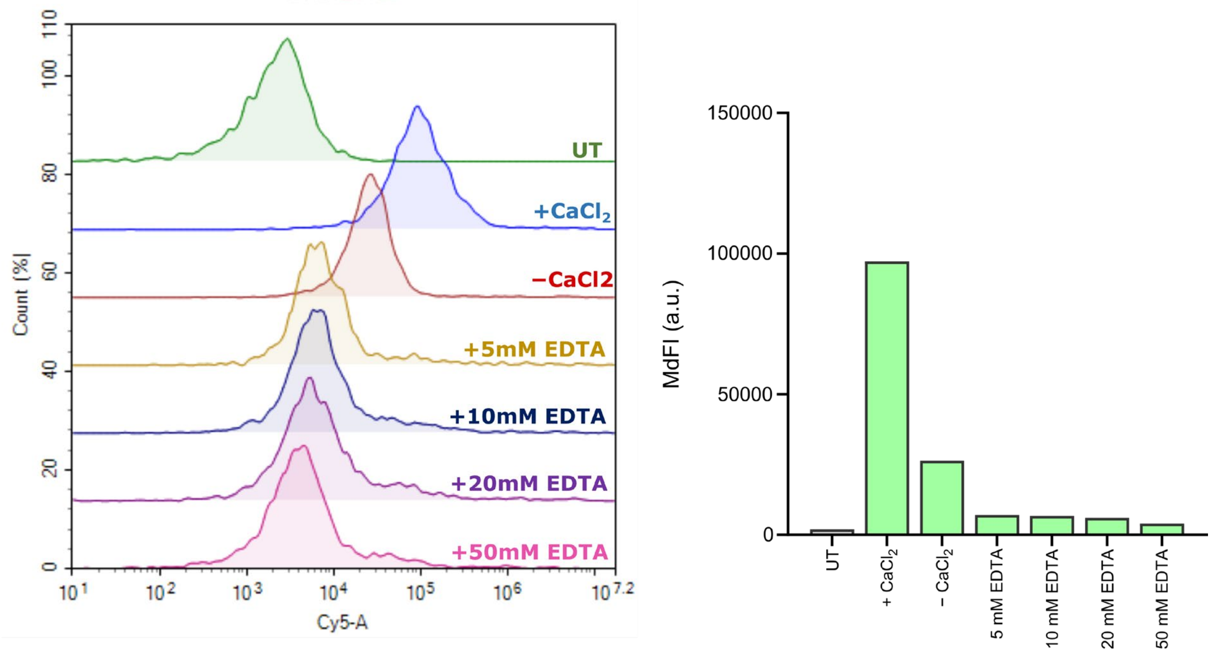


**Supplementary Figure S4. Removal of surface-bound mannosylated HJs from Langerin-expressing HEK293 cells.** The HEK293-Langerin+ cells were incubated at 4°C for 1 h in the dark with 50 nM Cy5-labeled HJ-3xTriMan, followed by washing under different conditions: cell staining buffer (20 mM HEPES pH 7.4, 5 mM $MgCl_2$, 5 mM $CaCl_2$, 150 mM NaCl, 0.5% w/v BSA, denoted as '+$CaCl_2$'), a calcium-free HEPES buffer ('−$CaCl_2$), and HEPES buffer containing EDTA of 5, 10, 20 or 50 mM. After washing, the cells were analyzed by flow cytometry for Cy5 fluorescence (*left*) and their MdFI was quantified (*right*), demonstrating that EDTA removes a majority of transmembrane bound mannosylated HJs.

Supplementary Figure S5

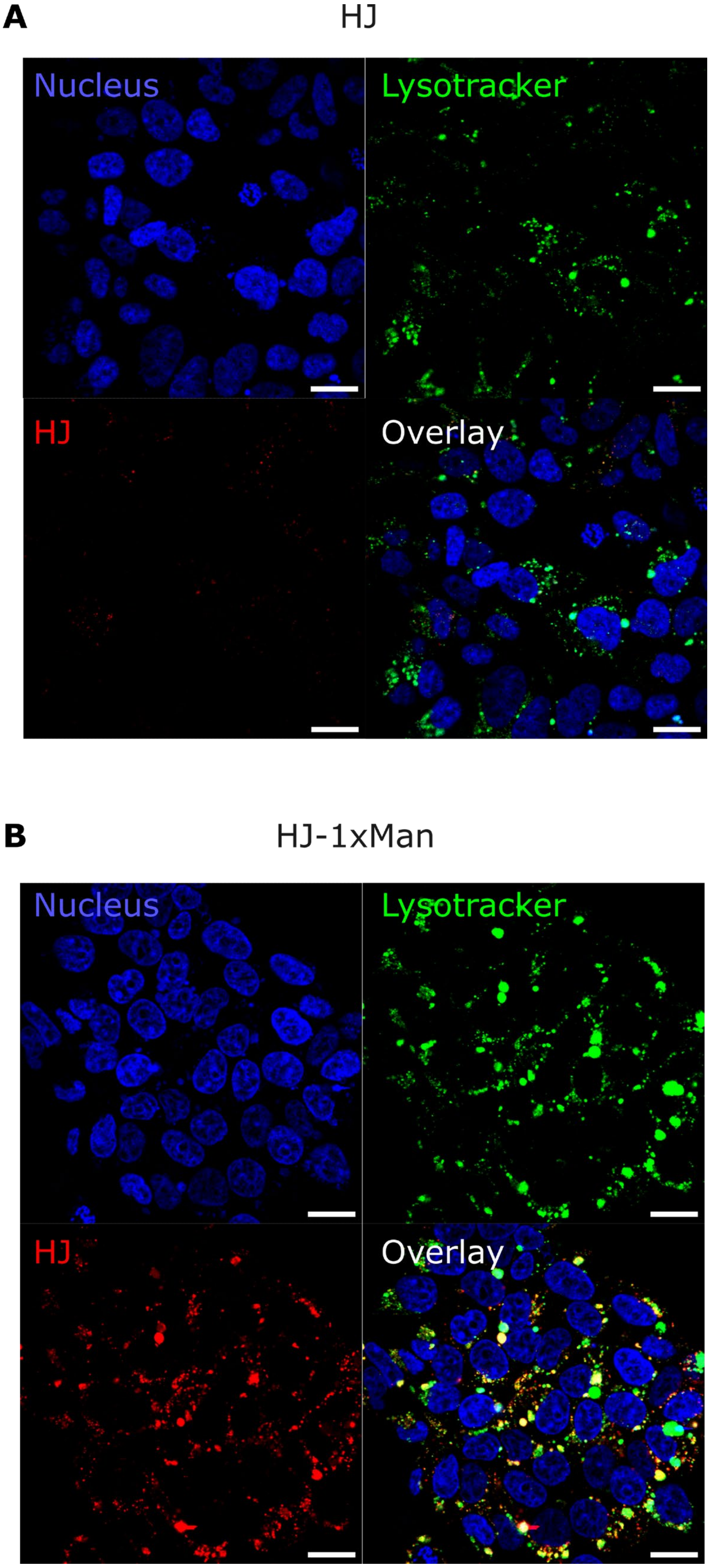

**C** HJ-2xMan

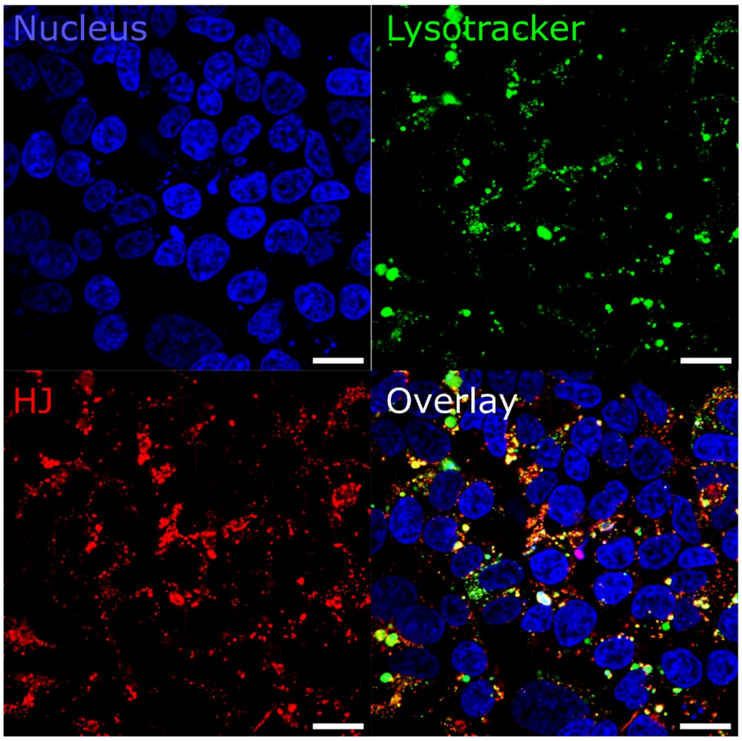


**D** HJ-3xMan

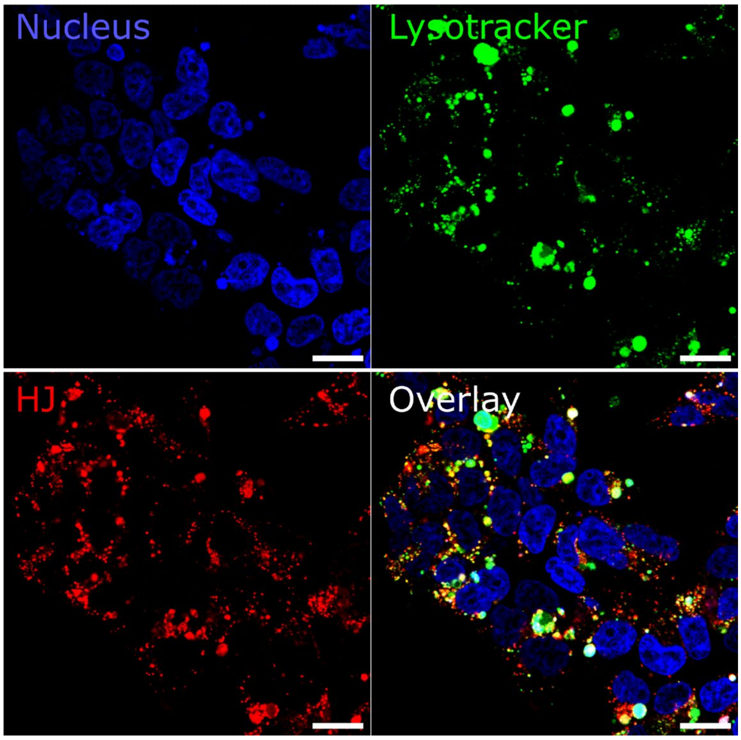

**E** HJ-1xTriMan

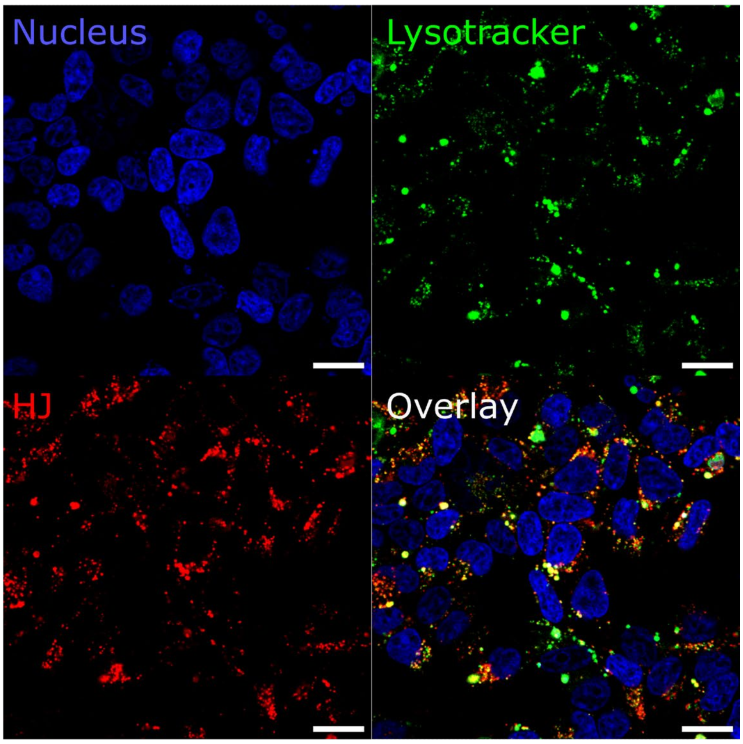


**F** HJ-2xTriMan

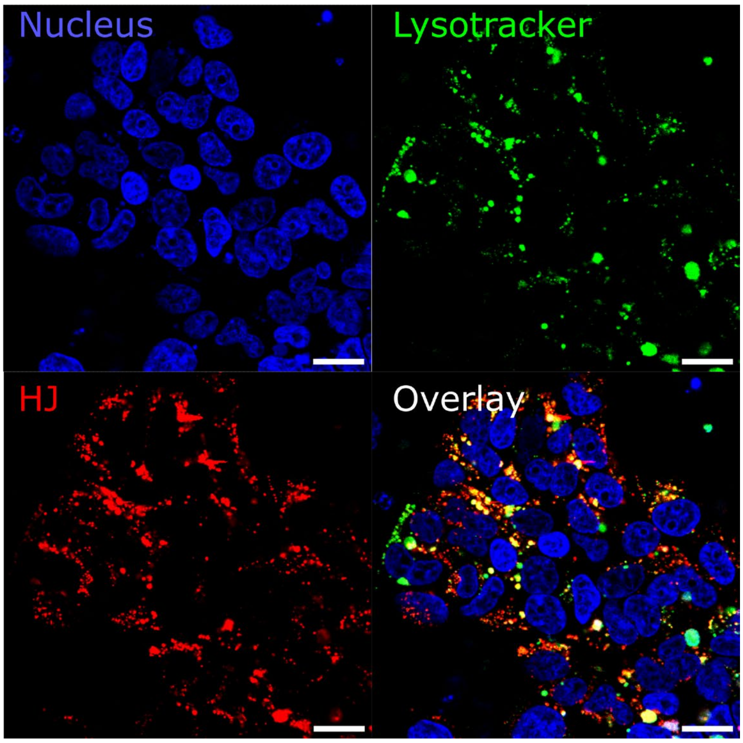

**G** HJ-3xTriMan

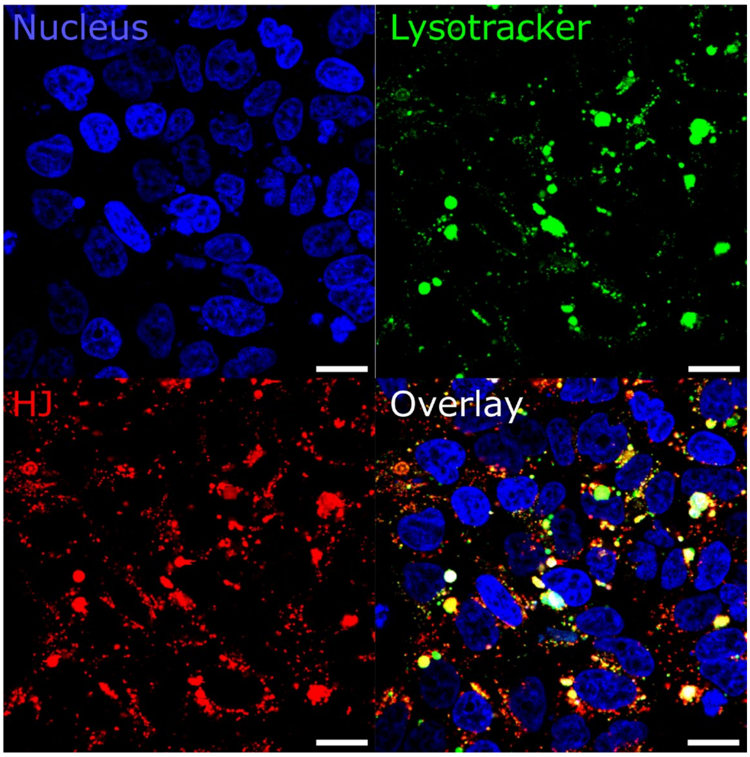


**Supplementary Figure S5. Full-sized confocal microscopy images of mannosylated HJs and lysosomal marker in Langerin-expressing HEK293 cells.** Live-cell confocal microscopy displaying cellular distribution of **A-G)** HJ, HJ-1xMan, HJ-2xMan, HJ-3xMan, HJ-1xTriMan, HJ-2xTriMan, HJ-3xTriMan treated HEK293-Langerin$^{+}$ cells. The HJ scaffolds were Cy5-labeled (red) and the cells were nucleus-stained with Hoechst 33342 (blue) and lysosome-stained with LysoTracker Green DND-26 (green). Lysosomal accumulation of HJs can be observed in the merged overlay of all three fluorescent channels (white). Scale bar = 20 µm.

#### Supplementary Figure S6

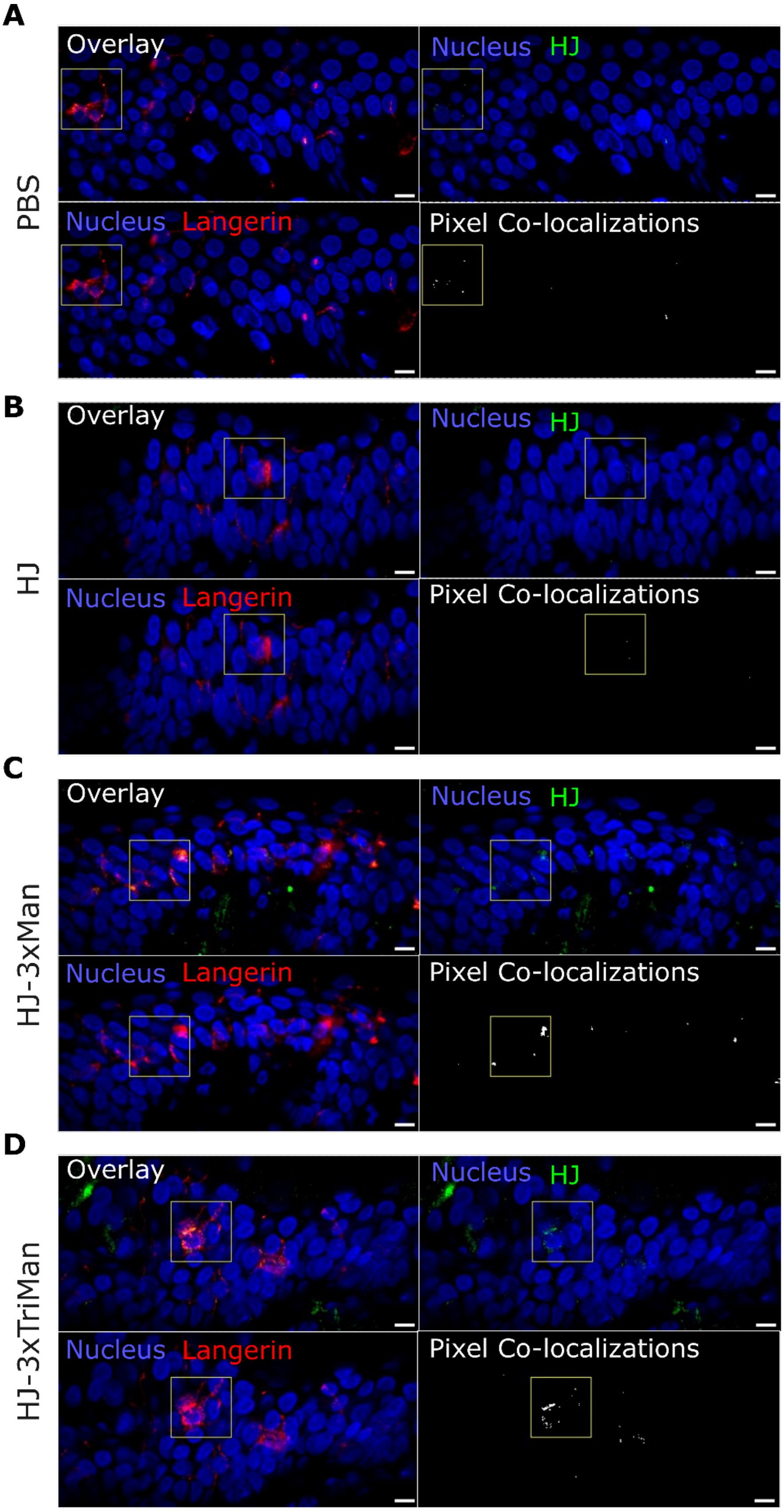

**Figure S6. Topical delivery of HJ onto Dermarolled human skin samples**. Representative images of skin samples subjected to topical delivery of PBS (A) and ATTO565-labeled HJ (B), HJ-3xMan (C) and HJ-3xTriMan (D), and visualized by confocal microscopy after nucleus-stained with DAPI (blue), Langerhans cells-stained with Langerin (red). Colocalization was conducted for the Langerin- and HJ fluorescent channel by pixel-based colocalization. The region of interest (yellow) enhanced in Figure 5 is shown with a yellow square. Scale bar = 20 µm.

### Supplementary Figure S7

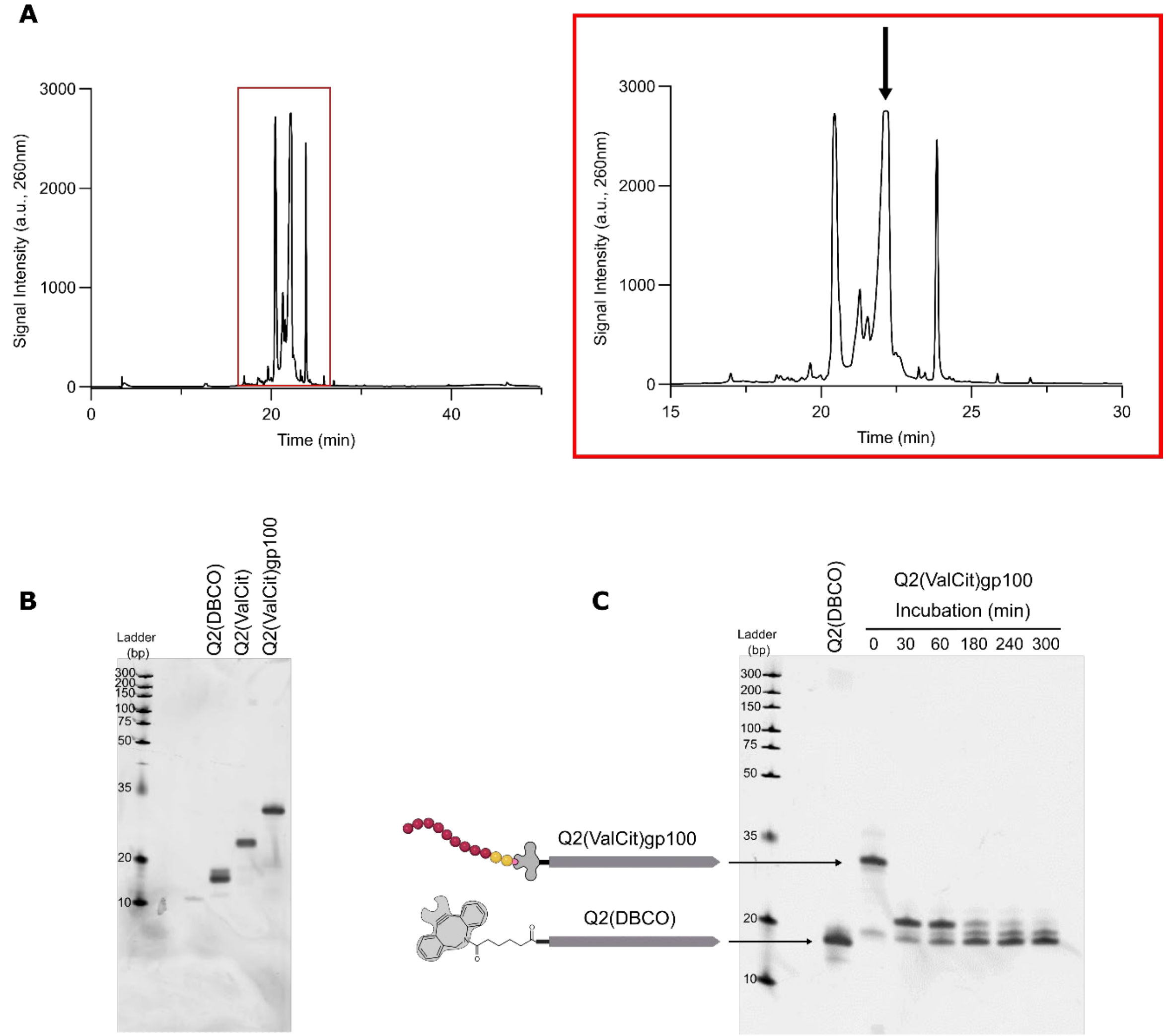


**Supplementary Figure S7. Purification and characterization of gp100-conjugation via a cleavable linker**. **A)** RP-HPLC purification of DBCO-functionalized Q2 reacted with azide-ValCit-PAB-gp100. The middle peak (arrow) in the chromatogram (Rt: 22.0 min) indicates the reaction product Q2(ValCit)gp100. **B)** The HPLC-purified Q2(ValCit)gp100 was analyzed on a 10% denaturing PAGE stained with SYBR Gold and compared to DBCO-functionalized Q2 (Q2DBCO) and azide-ValCit-PAB-PNP-functionalized Q2 (Q2ValCit). **C)** Cathepsin B-mediated cleavage of Q2(ValCit)gp100 after incubation for 0 – 300 min, visualized on a 12% denaturing PAGE stained with SYBR Gold. The release of gp100 by Cathepsin B was visualized by the lower migration of Q2(ValCit)gp100 after enzyme incubation, indicating cleavage of the Val-Cit linker.

### Supplementary Figure S8

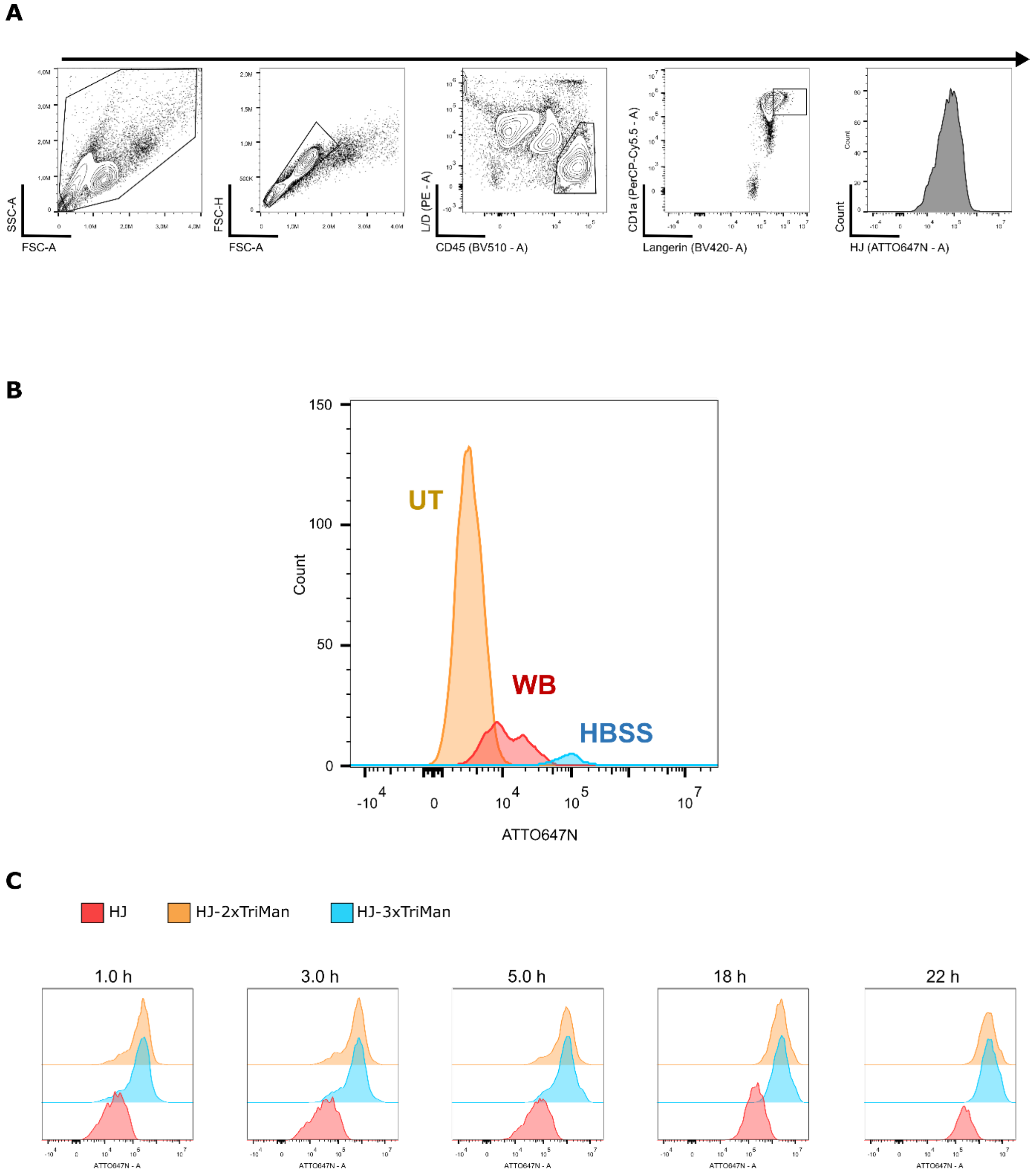


**Supplementary Figure S8. Kinetic uptake of HJ scaffolds into moLCs. A)** Flow cytometry gating strategy for moLCs ($CD45^{+}$, $CD1a^{+}$, $Langerin^{+}$) after 3 days of differentiation on a monolayer of OP9-$DLL4^{+}$ cells. **B)** MoLCs were incubated with 100 nM ATTO647N-labeled HJ-3xTriMan overnight at 4 °C to inhibit endocytosis, afterwards moLCs were washed with Hanks basic salt solution (HBSS, containing $Ca^{2+}$ and $Mg^{2+}$, blue) or an EDTA-containing WB buffer (red). Flow cytometry analysis of the moLCs revealed the majority of membrane-bound fluorescent HJ signal was removed after washing with WB buffer, comparable to untreated cells (UT, orange). **C)** MoLCs were incubated with 25 nM ATTO647N-labeled HJ (red), HJ-2xTriMan (orange) and HJ-3xTriMan (blue) for 1, 3, 5, 18 and 22 h at 37 °C, and washed with EDTA-supplemented WB buffer to remove membrane bound HJs, then analyzed by flow cytometry. The uptake of HJs was displayed by the ATTO647N fluorescence signal, visualized by their histograms.

### Supplementary Figure S9

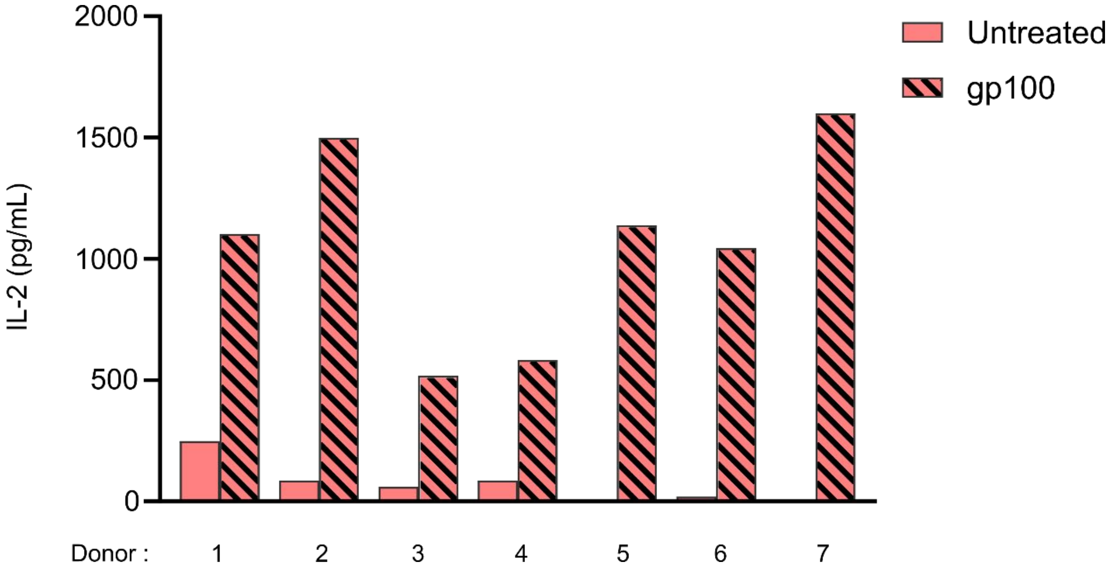


**Supplementary Figure S9. Collective positive controls for validation of antigen-presentation assay on moLC.** Bar graph of IL-2 secretion from gp100 Jurkat cells co-cultured with moLCs from seven different HLA-A2 positive donors loaded for 1 h with 2 µM $gp100_{154\text{-}162}$ antigenic peptide, compared to co-cultures with untreated moLCs. These data represent the positive controls gathered from the antigen presentation assay shown in Figure 6D-E.

### Supplementary Figure S10

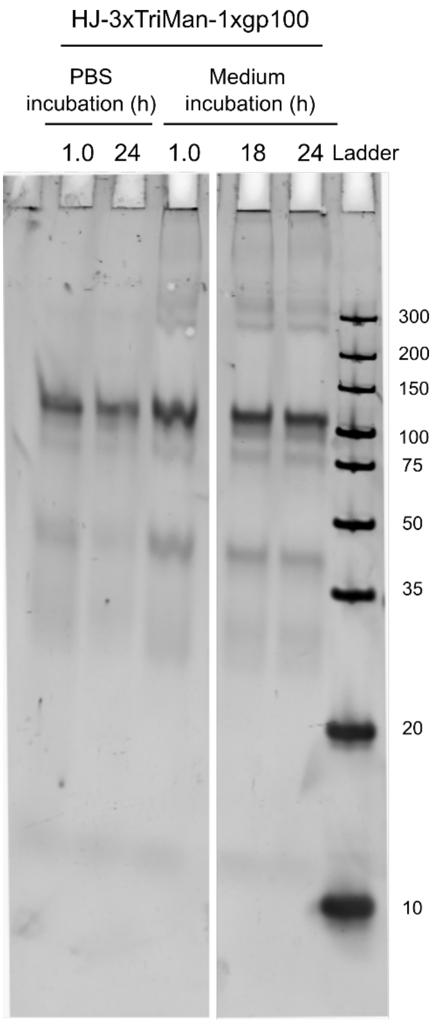


**Supplementary Figure S10. Medium stability of gp100-peptide conjugated HJ in R10 medium**. Stability of HJ-3xTriMan-1xgp100 in R10 medium, employed in the antigen-presentation assay, to assess the stability of the scaffold. HJ-3xTriMan-1xgp100 was incubated for 1, 18 and 24 h in R10 medium, and the sample was loaded on a 16% native PAGE stained with SYBR Gold to visualize scaffold integrity, compared to incubation for 1 and 24 h in 1xPBS.

### Supplementary Figure S11

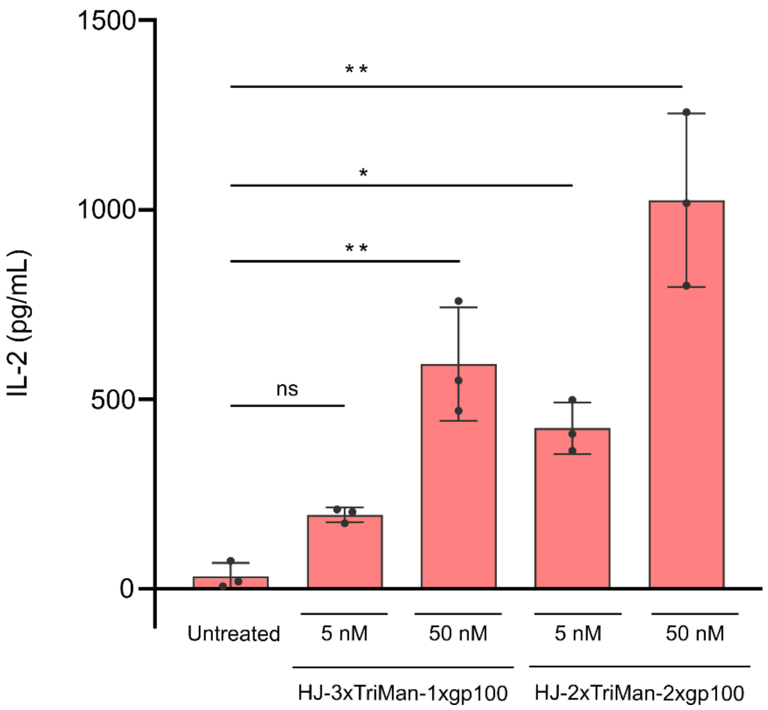


**Supplementary Figure S11. Antigen-presentation assay with mannosylated HJs in nanomolar concentrations.** HJ-3xTriMan-1xgp100 and HJ-2xTriMan-2xgp100 were used in an antigen presentation assay with moLCs at 50 and 5 nM concentrations. Results expressed as the mean ± SD for three different donors show a significant increase in IL-2 secretion at 50 nM for both scaffolds, however at 5 nM only the HJ-2xTriMan-2xgp100 was significantly increased when compared to the untreated moLCs. Statistical analysis was conducted by a one-way ANOVA followed by a Tukey's post hoc test. p-value * = <0.05, ** = <0.01, *** = <0.001, ****= <0.0001.
